# Repeated mutation of a GT92 glycosyltransferase gene confers antiviral resistance in two *Caenorhabditis* species

**DOI:** 10.64898/2026.04.14.718442

**Authors:** Aurélien Richaud, Gaotian Zhang, Cigdem Alkan, Daria Martynow, Tony Bélicard, Nanako Takeda, Eillen Tecle, Marie-Anne Félix

## Abstract

Host-pathogen interactions evolve rapidly within species, providing natural genetic resources for the identification of specific ecological interaction factors. We previously identified RNA viruses that infect the nematodes *C. elegans* and *C. briggsae* in a species-specific manner. Wild strains of both host species demonstrate ample variation in viral sensitivity. Specifically, the wild *C. elegans* strain MY10, despite carrying a deletion in a key immunity factor, was among the most resistant strains. Here we use recombinant inbred lines and pool-sequencing approaches to genetically map the major MY10 resistance locus, narrowing down its position by CRISPR/Cas9 mediated recombination and testing candidates by genome editing. A rare non-synonymous polymorphism in the *gtnt-1* gene, encoding a putative glycosyltransferase of the GT92 family, causes resistance to viral infection in MY10. We find that viral resistance through *gtnt-1* mutation occurred repeatedly in *C. elegans*, with diverse resistance alleles each remaining at low frequency (<1%). Furthermore, leveraging closely related *C. briggsae* strains differing in viral susceptibility, we demonstrate that repeated reduction-of-function alleles of the *Cbr-gtnt-1* ortholog similarly impair viral infection and enhance host fitness upon infection. In conclusion, we found recurrent evolution in two host species of reduction-of-function alleles of the *gtnt-1* orthologs, which repeatedly lead to viral resistance yet remain at low frequency. These repeated events provide a case of transient ecological adaptation to a pathogen through recurrent mutation of the same gene in two species. The low population frequencies of the resistant alleles point to a changing eco-evolutionary context that prevents their spread in populations, resulting in high allelic heterogeneity.

## Introduction

Do independent populations or species facing similar selective environments find similar molecular solutions? If so, this repeatability demonstrates that natural selection and constraints in the genotype-phenotype map channel molecular and phenotypic evolution despite the stochastic framework of mutation in which they operate (Martin and Orgogozo 2013). Pathogens exert strong and directional selection pressures on their hosts, making host-pathogen interactions ideal systems to study the repeatability of adaptive trajectories. A single host-pathogen pair allows asking whether the same pathogen drives parallel evolution independently across different host populations. Closely related host-pathogen pairs offer an even more powerful model. Although these hosts have diverged into distinct species with their own specificities, they share many orthologous genes. Likewise, their related pathogens may have conserved host dependency factors.

Hosts can respond to pathogens through two broad strategies: resistance via reduction of the viral load or tolerance to a high pathogen load. In host populations, reduction of the viral load can result either from individual resistance to infection or from a reduction in virion transmission across individuals. Therefore, experimental assays spanning several viral cycles after an initial inoculation are necessary to capture all aspects of the host-virus interaction, including propagation to other individuals in successive host generations.

Studying repeated host-pathogen adaptation and long-term interaction dynamics requires fast-growing, widespread host organisms with known natural pathogens. Selfing *Caenorhabditis* nematodes, *C. elegans* and *C. briggsae*, fulfill these requirements exceptionally well, with a lifecycle of only 3-4 days and large collections of wild isolates from across the globe. Crucially, through studies of natural populations of *Caenorhabditis*, a virus called the Orsay virus (ORV) was found to naturally infect *C. elegans*, and three related viruses, called Santeuil (SANTV), Le Blanc (LEBV) and Melnik (MELV) viruses were found to infect *C. briggsae*, with species specificity (Félix et al. 2011; Franz et al. 2012; Frézal et al. 2019). These non-enveloped RNA viruses are transmitted horizontally through the fecal-oral route and only infect host intestinal cells (Félix et al. 2011; Franz et al. 2014; Félix and Wang 2019). Their genome consists of two positive-strand RNA molecules. RNA1 codes for an RNA-dependent RNA polymerase (RdRp) and RNA2 for a capsid, related to those coded by viruses in the *Nodaviridae* family (Félix et al. 2011). Infection slows down host progeny production but does not strongly affect it, allowing viral propagation in reproducing host populations (Félix et al. 2011; Ashe et al. 2013; Alkan et al. 2024).

*C. elegans* and *C. briggsae* reproduce through selfing hermaphrodites and facultative males. Combined with a short lifecycle and the ability of these species to be cryopreserved, this mode of reproduction makes genetic approaches easy and powerful. Their interactions with natural viruses have thus provided tractable genetic models to discover antiviral immunity mechanisms and host factors required for the viral cycle (Félix and Wang 2019; Gonzalez and Félix 2024). In addition to laboratory genetic screens, wild strains of both species are natural genetic resources where polymorphisms affecting host-virus interactions can be mined. Genetic diversity of wild host strains can be used in two ways (Andersen and Rockman 2022): 1) genome-wide association studies (GWAS) that leverage recombination in natural populations, where a panel of strains is tested for statistical association between the phenotype of interest and polymorphic markers along the genome; 2) controlled laboratory crosses (typically between two strains of contrasted phenotype) and linkage mapping of recombinants along the genome.

In a previous study, we used the GWAS approach by assaying a panel of 97 *C. elegans* wild strains (Andersen et al. 2012) for their ability to propagate the Orsay virus (Ashe et al. 2013). To do so, we defined the phenotype as the viral load 7 days after viral inoculation, i.e. after two passages to new plates at 20°C. We detected an association on chromosome IV, explained by a causal polymorphism in the *drh-1* gene. We confirmed that a partial *drh-1* deletion allele caused the wild *C. elegans* strain JU1580 to accumulate a higher viral load compared to the more resistant N2 reference strain carrying an intact *drh-1* gene (Ashe et al. 2013). The DRH-1/RIG-I homolog triggers viral RNA degradation via small RNA pathways in *C. elegans* (Ashe et al. 2013; Guo and Lu 2013), as well as a host transcriptional defense response (Sarkies et al. 2013; Bakowski et al. 2014; Chen et al. 2017; Sowa et al. 2020). The *drh-1* deletion allele was found in 22/97 (23%) strains in the strain panel (Ashe et al. 2013). However, this polymorphism did not explain all of the phenotypic variation in this panel. Most strikingly, the MY10 strain carries the *drh-1* deletion but was among the strains displaying the lowest viral load in our assay (Ashe et al. 2013) (Figure 1a). In a more recent study, we surveyed 40 wild *C. briggsae* strains for infection rate by SANTV and LEBV using fluorescence *in situ* hybridization (FISH) at 7 days post-inoculation and revealed substantial variation (Alkan et al. 2024). Classical linkage mapping in RILs derived from a cross between AF16 and HK104 identified a major QTL on chromosome IV associated with variation in SANTV infection. In parallel, a bulk segregant pool-sequencing approach applied to a cross between JU1498 and HK104 identified a major locus on chromosome II underlying variation in LEBV infection.

**Figure 1.**
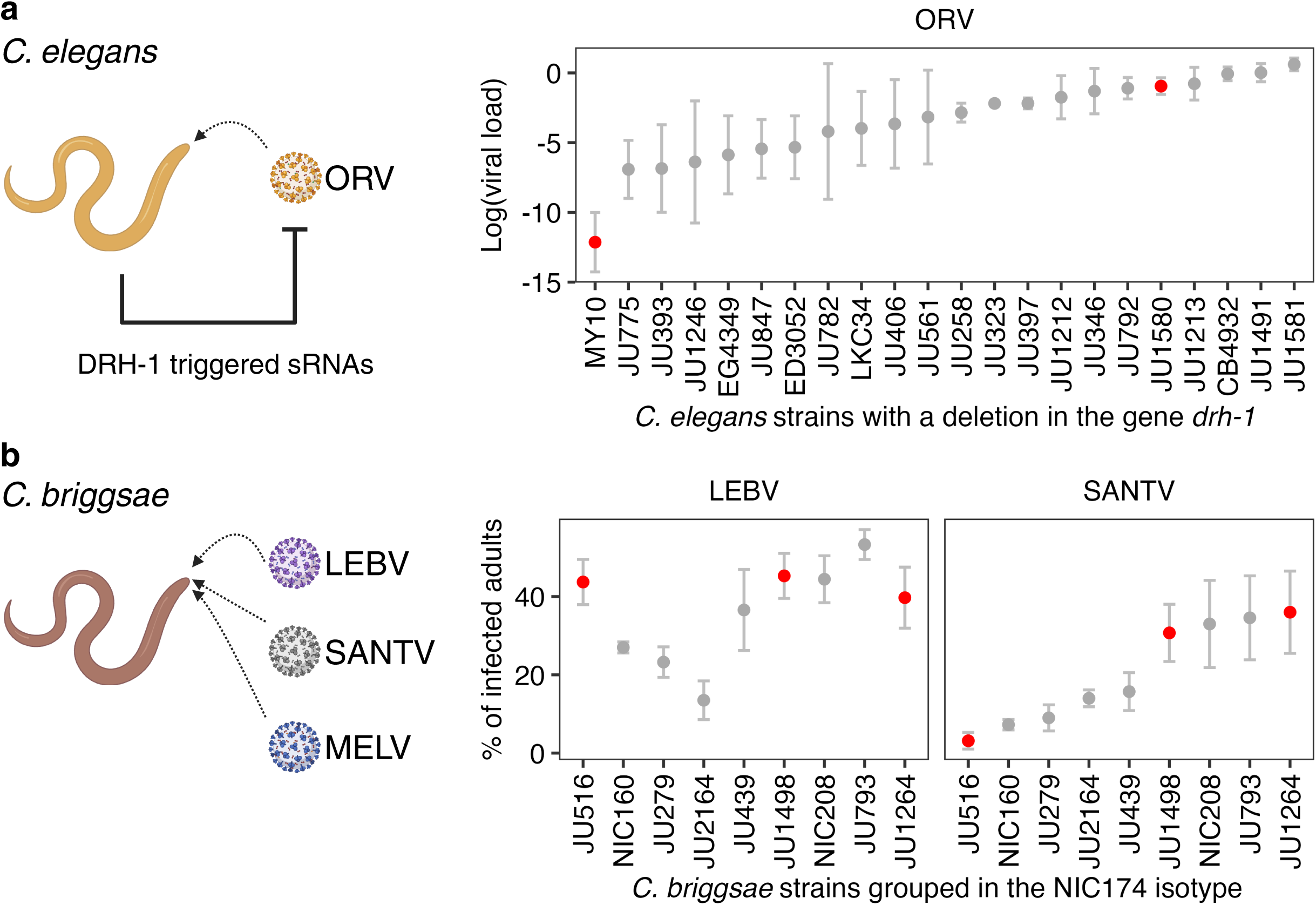
Natural variation in susceptibility of *C. elegans* and *C. briggsae* to viral infection. **a,** *C. elegans* can be infected by the Orsay virus (ORV), and uses small RNAs (sRNAs) as antiviral defense. The natural *drh-1(niDf250)* deletion prevents this antiviral sRNA response. The graph depicts variation in ORV replication across wild *C. elegans* strains carrying this deletion. Viral load (*y*-axis) at 7 days post-infection (dpi) was measured by qRT-PCR (Ashe et al. 2013). **b,** *C. briggsae* can be infected by LEBV, SANTV and MELV. Variation in infection rates of *C. briggsae* strains of the isotype NIC174 exposed to LEBV or SANTV. The percentage of infected individuals (y-axis) at 7-8 dpi was quantified by FISH (Alkan et al. 2024). Each point represents one strain, with error bars showing standard errors among replicates. The infection data in this figure are reproduced from prior publications (Ashe et al. 2013; Alkan et al. 2024) to highlight the rationale of the present study. Red dots mark strains selected for further analysis in this study.

Here we follow on two prior findings in our laboratory in these two *Caenorhabditis* species. First, we investigated the paradoxical finding that the *C. elegans* wild strain MY10, despite carrying the *drh-1* deletion, was among the strains with the lowest viral load in our assay (Figure 1a, modified from (Ashe et al. 2013). We used laboratory crosses to address the cause for MY10 viral resistance. After crosses of the MY10 genomic background to that of either the JU1580 or the JU1395 wild strains, we found that a locus on chromosome V explained the low infection rate of MY10 by the Orsay virus. We further mapped the locus using targeted CRISPR-Cas9-mediated recombination and found that a rare non-synonymous polymorphism (P182L) in the gene *gtnt-1* caused viral resistance of the MY10 strain. Other *C. elegans* wild strains with nonsense mutations or deletions in this gene also displayed ORV resistance, suggesting repeated evolution of viral resistance. Second, leveraging the recent high-throughput sequencing of *C. briggsae* strains (Crombie et al. 2024), we found that strains with different viral sensitivities in our dataset (Figure 1b) (Alkan et al. 2024) had very similar genomes, which led us to find independently arising alleles of the *gtnt-1* ortholog in *C. briggsae.* Taken together, reduction-of-function alleles of the *gtnt-1* orthologs are found at low frequency in both species, thus demonstrating repeated molecular evolution leading to parallel evolution of viral resistance.

## Methods

### Culture and strains

*C. elegans* and *C. briggsae* were grown under standard conditions (Stiernagle 2006). A bleach treatment was applied to all isolates prior to virus infection to eliminate possible contaminations, as in (Stiernagle 2006; Félix et al. 2011). Viral filtrates of the ORV strain JUv1580, the SANTV strain JUv1264, the LEBV strain JUv1498, and the MELV strain JUv3272 (Frézal et al. 2019) were prepared as described previously (Félix et al. 2011). A list of *Caenorhabditis* strains used in this study can be found in Table S1.

### Introgression of the *lys-3p::GFP* transgene in JU1580 and MY10 backgrounds

The *mjIs228[myo-2p::mCherry::unc-54; lys-3p::eGFP::tbb-2]* transgene from SX2790 carrying a reporter of viral infection in the *C. elegans* N2 reference background (Le Pen et al. 2018))was introgressed into the MY10 and JU1580 genomic backgrounds through 10 backcrosses, following the transgene through the red fluorescence in the pharynx conferred by the *myo-2p::GFP* transgene. The choice of these two strains derives from previous results shown in Figure 1a (Félix et al. 2011; Ashe et al. 2013). The introgressions yielded the strain JU2624 (JU1580 background) and JU2625 (MY10 background). We mapped the transgene to chromosome II using the mapping strains EG1000 and EG1200 obtained from the Caenorhabditis Genetics Center.

### Viral infection assays on *C. elegans*

Our assays were designed to account for all stages of the viral cycle over at least two host generations. Before viral inoculation, 10 L4 stage larvae of a previously bleached culture were placed onto 55 mm NGM plates seeded with *E. coli* OP50. 30-40 μL of filtrates of the viruses were added into the center of the *E. coli* OP50 lawn. Inoculated cultures were incubated at 20°C for 5-7 days. Maintenance of the infected cultures was performed by transferring a piece of agar every 2-3 days to a new plate with food but without additional viral filtrates.

In some experiments in Figure 3, bleached embryos were deposited on the plate and the viral preparation was added, thus first exposing L1 larvae.

### Fluorescent *in situ* hybridization (FISH) for viral RNA

The protocol is adapted from (Franz et al. 2014) with the multiple RNA2 probes used in (Franz et al. 2014; Frézal et al. 2019). Nematodes were harvested with nuclease-free water and pelleted by centrifugation at 3000 rpm for 5 min. The pellet was transferred to a non-adhesive 1.5 mL tube (Axygen). The fixative solution contained 5 mL 37% formaldehyde (Sigma #533998), 5 mL 10x PBS (Ambion AM962, pH 7.4), 40 mL nuclease-free water. 1 mL of this solution was added to each tube, and the tube was rotated with light agitation for 40 min. After removing the fixative solution, the pellet was washed twice with 1x PBS (Invitrogen), resuspended in 70% ethanol and stored overnight at 4°C. The animals were pelleted and resuspended with the wash solution - 5 mL deionized formamide (Ambion AM9342) (10 mL for *C. briggsae*), 5 mL 20X SSC (Ambion AM9770) and adding nuclease-free water to 50 mL. They were resuspended in 100 μl hybridization buffer. The hybridization buffer contained for 10 mL: 1 g dextran sulfate (Sigma #D6001), 1 mL 20x SSC, 1 mL deionized formamide (2 mL for *C. briggsae*), and adding nuclease-free water to 10 mL. 1 µL of 1:40 diluted fluorescent probes were added and the fixed animals were incubated overnight at 30°C in the dark. The following day, they were washed with 1 mL wash solution, resuspended in 0.02% DAPI (Sigma #D9564, 5 mg/mL) in 1 mL wash solution and incubated for 30 min at 30°C in the dark. Finally, they were resuspended in 2x SSC and could be kept at 4°C before imaging. For imaging, ∼4 µL of sample were pipetted onto round coverslips, which were sealed onto glass slides using silicon isolators. The fluorescence of adult animals was scored with an Olympus FV1000 macroscope or a Zeiss AxioImager M1.

### Recombinant Inbred Lines (RILs)

Recombinant inbred lines were built by crossing the two parental lines, isolating single F2 progeny and selfing them for 10 more generations. Because of the strong temperature-dependent mortal germline phenotype of the MY10 strain (Frézal et al. 2018), the two parental lines were crossed at 15°C and F2 cross-progeny individuals were then selfed for 10 generations at 15°-18°C (Figure 2b).

**Figure 2.**
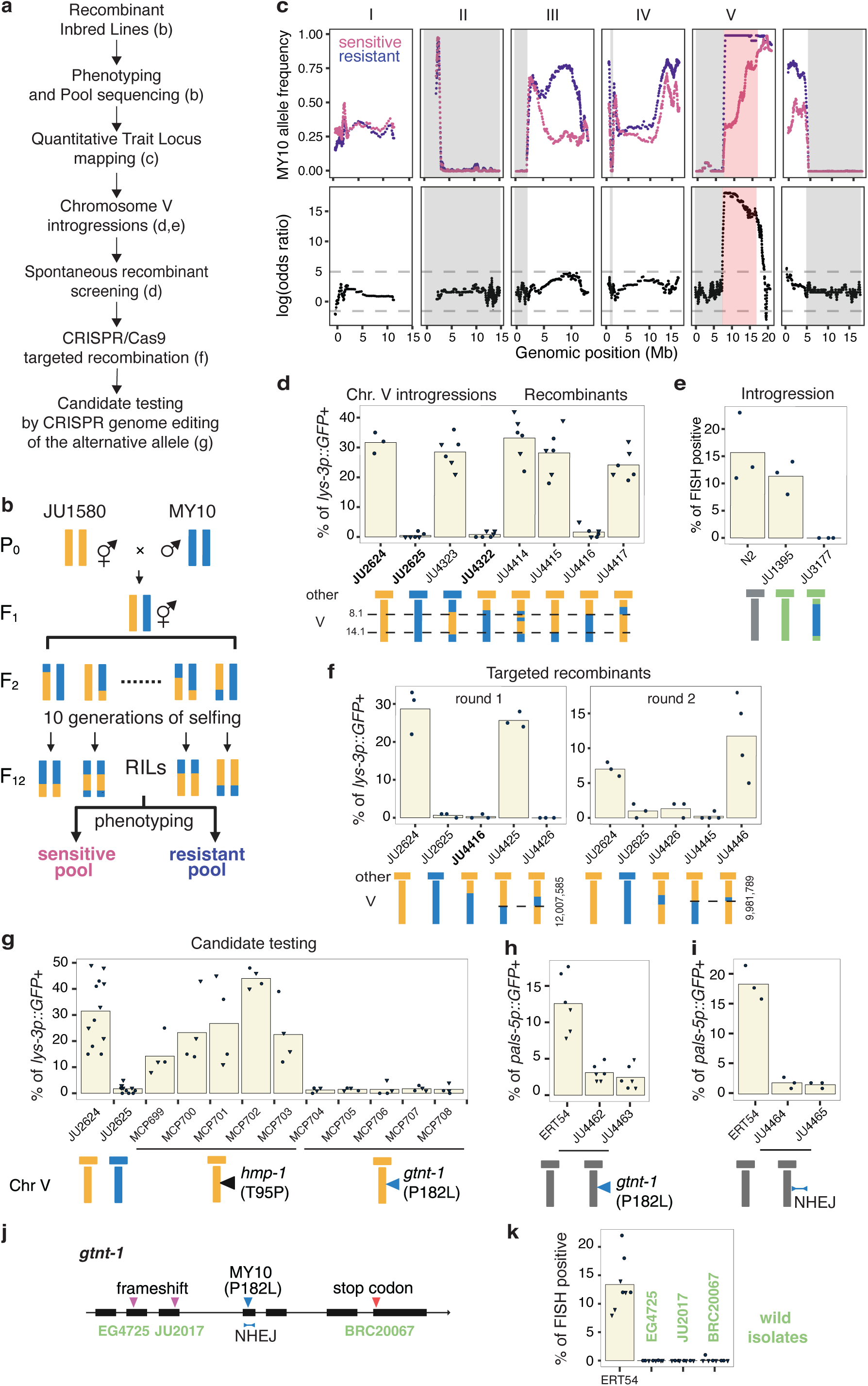
Genetic dissection of viral resistance in the *C. elegans* strain MY10 identifies a causal *gtnt-1* polymorphism. **a,** Overall design of the genetic dissection of viral resistance in the *C. elegans* strain MY10, with reference to corresponding figure panels. **b,** Crossing scheme used to generate recombinant inbred lines (RILs) between the MY10 and JU1580 wild backgrounds. RILs were phenotyped for ORV replication by FISH at 7-8 dpi, identifying 27 resistant and 26 sensitive lines. **c,** Bulk sequencing of pooled resistant and sensitive RILs. The MY10 allele frequency of each pool was plotted along the *C. elegans* genome, with corresponding LOD scores shown below. Dotted lines represent the significance threshold (*p*<0.01). Grey shades represent regions without polymorphism among RILs (see Methods). Pink shade represents the QTL region of interest. **d,** Reciprocal chromosome V introgressions confirm the QTL and spontaneous recombinants restrict the interval to 8.1-14.1 Mb on chromosome V. The genotype of each strain is schematized below the graph, the vertical bar denoting chromosome V and the horizontal bar the rest of the genome. Orange: JU1580; blue: MY10. JU2624 and JU2625 are JU1580 and MY10 derivatives with the *lys-3p::GFP* reporter (not shown in the schematic). Infection was scored using this reporter at 5 dpi after inoculation of 10 L4 larvae on 3 replicate plates in two independent experimental blocks, indicated by datapoint shapes. **e,** Chromosome V introgression from MY10 into JU1395 (Frézal et al. 2018) also leads to resistance to ORV infection, as scored by FISH against the virus at 4 dpi in 3 replicate infections. Grey: N2; green: JU1395. **f,** CRISPR-Cas9 induced recombination at locations indicated by the two dashed lines narrowed down the interval to 9.982-12.008 Mb. For each experiment, 3 replicate plates with 10 L4 larvae each were inoculated with ORV and the animals were scored at 5 dpi. 100 animals were scored per replicate. **g,** Replacement in the JU1580 background carrying the *lys-3p::GFP* reporter of two candidate genes (*hmp-1* and *gtnt-1*), edited to match their MY10 allele. Infection rate was scored using activation of the reporter at 5 days post-inoculation (dpi), in two independent experimental blocks indicated by datapoint shape. **h,** Replacement of *gtnt-1(P182L)* in the N2 background with the *pals-5p::GFP* reporter prevents reporter activation upon ORV inoculation. **i,** Insertion-deletion *gtnt-1* alleles obtained by non-homologous end joining (NHEJ) in the N2 background with the *pals-5p::GFP* reporter prevents reporter activation upon ORV inoculation. JU4464 carries an inframe indel and JU4465 carries a frameshift (Table S1). **j,** *C. elegans gtnt-1* gene structure, with positions of the MY10 missense, the NHEJ lesions and the tested natural variants. **k,** Wild *C. elegans* strains carrying stop-gain or frameshift mutations (**j**) were tested for viral resistance using FISH. See Table S4 for raw data.

To assay ORV infection over the full viral cycle, the lines were inoculated with an ORV JUv1580 filtrate and fixed 7-8 days later. The lines were then scored by FISH against viral RNA2, each twice in independent inoculations (two blocks started on different days).

Genomic DNA of 27 lines that were consistently negative in both blocks were pooled to form the “resistant pool”. Genomic DNA of 26 lines that were clearly positive (at least 20% FISH-positive animals) in at least one block were pooled to form the “sensitive pool”.

After genomic sequencing of the pools (see below), we realized that two errors (likely related) had been made in the construction of the RILs. First, the *lys-3p::GFP* transgene was not present in all the RILs thus was likely not homozygous in the parents at the start of the RILs. Hence it was not used for scoring the RILs. The transgene insertion was homozygous in the JU2624 and JU2625 introgressed strains and was later used in the fine mapping (see below). Second, some large regions of the genome, for example on the left of chromosome V (Figure 2c) appeared homozygous for the JU1580 allele in both pools, with a boundary at the same nucleotide position in both pools (for example at nucleotide 7675199 on chromosome V). A possibility is that one of the backcrosses of the *lys-3p::GFP* to the MY10 strain was erroneously performed by crossing it to JU1580 instead. The JU2625 strain that we froze is however in the MY10 background except for the chromosome II introgression. Alternatively, the cross for the RILs was performed using as “JU2625 mothers” the product of a prior cross (the cross had to be performed twice because of high sterility at 20°C; the second cross was performed at 15°C). The QTL mapping is thus only informative for the part of the genome in which both MY10 and JU1580 alleles were present.

### Pooled genome sequencing and linkage mapping

The design was similar to that previously used in (Frézal et al. 2018; Dubois and Felix 2023) and for the second set of RILs in (Alkan et al. 2024). Specifically, the genomes of the two pools of resistant and sensitive RILs were sequenced using paired-end Illumina sequencing. Genomic DNA was extracted from mixed-stage populations of each line using the Puregene Core Kit A (QIAGEN, Valencia, CA) and concentrations measured with Nanodrop (Thermofisher) and adjusted in the pool to provide equal representation of each line. Paired-end sequencing using Illumina HiSeq 4000 at 30x coverage was performed by BGI genomics with paired-end 100 bp reads. Reads are available at NCBI with accession number PRJNA1435841.

In the analysis shown in Figure 2c, adapter sequences and low-quality reads in raw sequencing data of the two pools were removed using *fastp* (v0.20.0) (Chen et al. 2018). The trimmed reads were aligned to the six chromosomes (I, II, III, IV, V, X) of the *C. elegans* N2 reference genome (Wormbase WS283) (Davis et al. 2022) using *BWA* (Li and Durbin 2009) incorporated in the pipeline *alignment-nf* (https://github.com/AndersenLab/alignment-nf) (Cook et al. 2017). Whole-genome alignments of the two parental strains, MY10 and JU1580, were obtained from CaeNDR (20231213 release). Single nucleotide variant (SNVs) among the two parental stains and the two pools were called against the reference using *GATK* (v4.1.4) (Poplin et al. 2018) incorporated in the pipeline *wi-gatk* (https://github.com/AndersenLab/wi-gatk/) (Cook et al. 2017).

We performed a bulk segregant analysis to detect genomic regions where parental allele frequencies deviate between the ORV sensitive and resistant pools. We calculated the frequency of the MY10 allele at each SNP for each pool and filtered out SNPs in the MY10 and JU1580 hyper-variable genomic regions, where calls are uncertain (CaeNDR 20231213 release; (Crombie et al. 2024)). A total of 46,831 SNPs were used in the following analysis (Table S3). Then, we calculated mean frequency per 10 kb bin in the genome for each pool. We further performed a sliding window analysis with a 25-markers (binned SNPs) window size and a one-marker step size.

To test whether MY10 allele frequencies significantly differed between the sensitive and resistant pools from the expectation under a random distribution, we calculated for each window the log-odds ratio as: *log(m_1_/(n_1_-m_1_))/ (m_2_/(n_2_-m_2_)))*, *m_1_* and *m_2_* being the MY10 allele frequency multiplied by the number of RILs in the sensitive (n_1_=26) and resistant pools (n_2_=27), respectively. The significance thresholds at *p* = 0.01 were estimated by simulating the log-odds ratios using the binomial distribution across one million randomized draws given the number of strains in the two pools, as in (Frézal et al. 2018).

To find molecular markers for genotyping, the Pindel software (Ye et al. 2009) was used to detect homozygous indels in the JU1580 and MY10 strains. Parent-specific deletions were identified and manually checked using Tablet (Milne et al. 2013).

### Chromosome V introgressions and fine mapping using Near Isogenic Lines (NILs)

The introgressed lines JU4322 and JU4323 were built to confirm the QTL region detected through the bulk segregant analysis. We crossed JU2625 L4 hermaphrodites with JU2624 males. The male F1 cross progeny were then crossed to either JU2624 or JU2625 hermaphrodites, to introduce chromosome V of JU1580 and MY10, respectively, in the other background, following a single-nucleotide polymorphism (SNP) at the position V: 7,854,705 (G/C polymorphism in *anr-36* non coding RNA) using pyrosequencing using a PyroMark Q96 ID instrument (Biotage), according to the manufacturer’s instructions. The list of oligonucleotides is in Table S2. The backcross was repeated during 6 rounds.

Further NILs were obtained through random recombination at meiosis or by biasing the recombination locus by CRISPR (Zdraljevic et al. 2023). To generate recombinants, we crossed JU4322 hermaphrodites to JU2624 males and screened with pairs of primers AR5-6 and AR43-44 (Table S2). The four recombinants obtained (JU4414, JU4415, JU4416 and JU4417) were then genotyped by PCR or pyrosequencing with markers between 6.7 and 18.4 Mb on chromosome V using primers listed in Table S2. Further recombinants JU4425, JU4426, JU4445 and JU4446 were generated by inducing recombination at specific position V:9,981,789 and V:12,007,585 according to the method described by (Zdraljevic et al. 2023). Briefly, we crossed JU4416 L4 hermaphrodites with JU2624 males. The F1 heterozygotes were injected at the young adult stage with gRNA_4 (12,007,585) or gRNA_6 (9,981,789) mixed with a Cas9 ribonucleoprotein injection mixture and a *dpy-10* co-injection marker (Arribere et al. 2014). The F2 progenies of the injected animals that expressed the Rol or Dpy phenotypes were singled on plates and screened for recombination with primer pairs AR7-8 and AR84-85

### Caenorhabditis genome editing

The initial Cas9-mediated genome editing of several candidates in the *C. elegans* JU2624 background was performed by the SegiCel platform of CNRS in Lyon, France. Replacement was performed without altering the recognition site else than by introducing the desired nucleotide change. We detail below the method using the *gtnt-1* gene as an example. Editing of this gene in *C. elegans* backgrounds other than JU2624 was performed by us.

The repair template oMG286 for targeted replacement in the *gtnt-1* gene was designed to replace the C nucleotide at position V:10111167 in ERT054 to a T as in MY10, which results in a Proline to Leucine amino acid change (see sequence in Table S1). Before injection, 2 μL of 200 μM guide RNA crMG079 (synthesized by IDT) and 1.5 μL of 200 μM tracer RNA (IDT) were incubated in a PCR machine at 95°C for 3 minutes followed by decreasing temperature steps of 5°C steps every minute until 25°C. We used a guide RNA targeting the *Cbr-dpy-10* gene as a co-CRISPR marker. 1.0 μL of 200 μM *dpy-10* guide RNA and 0.7 μL of 200 μM tracer RNA were pre-incubated using the same temperature protocol. The final injection mix was: 3.5 μL of crRNA-tcRNA, 1.7 μL of dpy-10 crRNA-tcRNA, 0.3 μl Cas9 (IDT, 10 ng/μL), 1 μL of each repair template (Eurofins, 110ng/μL), 2.5 μL of nuclease free water (Life technologies). This mix was pre-incubated at 37°C for 30 minutes before injection.

Young adults were injected with the injection mix and transfered to new *E*. *coli* OP50 plates. After two days, if dumpy or roller progeny (co-CRISPR marker) was seen on the P0 plates, the plate was selected and several F1 progeny were singled. F2 progenies were screened by a PCR targeting the region around the edit with primer pairs AR96-97. 24 PCR products were sequenced by Sanger sequencing to find the correct replacement. Two strains harboring the replacement (JU4462 and JU4463) were kept. A second round of injection without repair template was performed to generate the two knock-out lines JU4464 and JU4465 by screening small deletions by PCR.

The same protocol was used in WUM32 background (Jiang et al. 2017) to generate the same SNP replacement (JU4632) and a knock-out deletion (JU4633).

The Cas9-mediated genome editing of *C. briggsae* MCP833 and MCP919 in the JU1498 and JU1564 background, respectively, were performed by the SegiCel platform.

All RNA oligonucleotides and PCR primers are listed in Table S2.

### *C. elegans* GFP viral infection reporter scoring

We used *pals-5p::GFP* (Bakowski et al. 2014) and *lys-3p::GFP* (Le Pen et al. 2018) as reporters for the infection. The animals were scored under a Leica MZ FLIII GFP stereomicroscope. Only adults were scored.

The WUM32 strain with the heat-shock construct expressing the ORV RNA1 as well as the *gtnt-1*edited lines were heat-shocked for one hour at 37°C and scored for *pals-5p::GFP* expression after 24 hours.

### Infection tests with the microsporidian parasite *Nematocida parisii*

*N. parisii* spores were isolated as previously described (Balla et al. 2015). 1500 synchronized L1 were grown to L4 and then were mixed with 3.2×10^6^ *N. parisii* spores, 10X concentrated OP50-1 bacteria (Wernet et al. 2025). This mixture was then plated onto room-temperature, unseeded 6 cm NGM plates and incubated at 25°C for 3 hours. The animals were washed twice in M9, plated on 1X OP50-1 seeded NGM plates, and grown for 21 hours at 25°C. Animals were fixed in 4% paraformaldehyde, stained using a FISH probe specific to *N. parisii* ribosomal RNA conjugated to Cal Fluor 610 dye (Biosearch Technologies), and imaged with a 10x objective on a Zeiss AxioImager M2 microscope. Using FIJI software, the integrated density of FISH probe fluorescence per worm was quantified, and background fluorescence was subtracted, yielding the Corrected Total Fluorescence (CTF) for each worm. CTF was normalized to the worm area. For one biological replicate, 60 animals per genotype were quantified, and three biological replicates were performed.

### Exposure to nematotoxic fungal lectins

Five fungal lectins were tested alongside a control plasmid pET24b: CGL2 (Butschi et al. 2010), LbTectonin2 (Wohlschlager et al. 2014), MOA (Wohlschlager et al. 2011) and CCTX2 (Cordara et al. 2011; Schmieder et al. 2025). 10ml of LB + Kanamycin (50mg/ml) were inoculated with a freshly transformed BL21 (DE3) and the cultures let grown at 37°C until OD600 reach between 0,5 and 1. Then, the protein expression was induced during 2 hours at 16°C by adding 0.5 mM of isopropyl-b-D-thiogalactoside (IPTG). 100 ml of these bacterial cultures were spread on 60 mm NGM-plates containing 1 mM of IPTG and incubated overnight at 23°C before addition of the nematodes. Five L4 stage hermaphrodites were isolated on five individual plates seeded with *E. coli* BL21 expressing the desired lectin as in (Butschi et al. 2010). The hermaphrodites were transferred each day to a new plate expressing the same lectin. Brood size was counted on every plate as the number of viable, motile larvae.

### Identification of polymorphisms within the *C. briggsae* NIC174 isotype

Short-read whole-genome sequencing alignments of the wild *C. briggsae* strains JU516, JU1498, and JU1264 to the reference strain QX1410, along with the reference genome, were obtained from CaeNDR (Andersen et al. 2012; Cook et al. 2017). SNVs and short indels were called against the reference genome as described above. Genetic variants were subsequently annotated using the *bcftools csq* function (Danecek et al. 2021) implemented in the pipeline *annotation-nf* (https://github.com/AndersenLab/annotation-nf).

### Multigenerational competition assay

Multigenerational competition assays among three *C. briggsae* strains (JU516, JU1264, and JU1498) were performed under four experimental conditions: uninfected control and infection with LEBV, SANTV, or MELV. All assays were conducted in parallel at 25°C. For each competition, five L4 larvae of each strain from a bleached culture were placed together on a 6 cm NGM plate seeded with *E. coli* OP50. After 24 h, 150 µL of either M9 solution (control) or a viral preparation was distributed evenly onto the bacterial lawn. Each condition included four biological replicates. On day 3, animals from each plate were collected by washing with 1.5 mL M9. From the well-mixed suspension, 100 µL was transferred to a fresh 10 cm NGM plate seeded with *E. coli* OP50 to generate two technical replicates per biological replicate. The remaining suspension was centrifuged, and 2 µL of the pellet was stored at −80°C for DNA extraction and subsequent pyrosequencing (Transfer 0). Populations on 10 cm plates were monitored daily. Prior to food depletion, 100 µL of a well-mixed suspension was transferred to a new 10 cm plate, and the remaining population was pelleted and stored as described above. Contaminated plates were discarded and replaced using independently maintained technical replicates. If both technical replicates for a biological replicate were contaminated, replacement plates were generated from other biological replicates of the same treatment. A minimum of four replicates was maintained for each condition throughout the experiment. The assay was carried out for 19 days and nine transfers (Transfers 0-8).

Multigenerational pairwise competition assays were further performed with strains JU1498 and its *CBG09519* edited derivative MCP833, each competing individually against JU516 in parallel, as described above. The pairwise assays were repeated once, with the first and the second experiments lasting 21 and 22 days, respectively. Samples from Transfer 0 and the final transfer (Transfer 9) were subjected to DNA extraction and pyrosequencing.

Strain frequencies in each sample were quantified by pyrosequencing allele-specific SNPs based on the reference strain QX1410: I:8995866 (A/G) to distinguish JU516 (G) from JU1264, JU1498 and MCP833, and V:9697991 (C/T) to distinguish JU1498 (T) from JU516 and JU1264, using previously described methods (Duveau and Félix 2012) with SNP-specific primers (Table S2).

### Estimation of relative fitness and selection coefficient

The relative fitness (*w*) of JU1498 and MCP833 compared to JU516 was estimated as previously described Zhao et al. 2018):

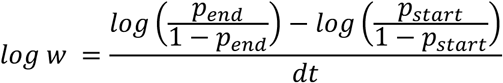

Where *p_start* and *p_end* denote strain frequencies at the beginning and end of the assay over *dt* generations. One generation per transfer was assumed in the competition assays. To avoid divergence of the odds ratio when allele frequencies approached 0, a value of 0.01 was used instead.

The selection coefficient *s* was then calculated from *w*:

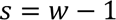

Statistical significance (*p*) and effect size (*r*) of differences in the selection coefficient *s* of JU1498 and MCP833 relative to JU516 was calculated using the Wilcoxon test via the functions *wilcox_test()* and *wilcox_effsize()*, respectively, in the R package *rstatix* (v0.7.2) (https://CRAN.R-project.org/package=rstatix). Resulting *p*-values were adjusted for multiple testing using the Benjamini-Hochberg false discovery rate (FDR) method via the base R function *p.adjust()*.

### Serial diluted viral infection assay on *C. briggsae*

Viral preparations of LEBV, SANTV, and MELV were successively diluted 3-fold to generate 1×, 1/3×, 1/9×, and 1/27× preparations. On Day 1, five L4 larvae from a bleached culture of a strain were placed on 6 cm NGM plates seeded with *E. coli* OP50 and maintained at 25°C. On Day 2, 150 µL of either undiluted or diluted virus was distributed evenly onto the bacterial lawn. Each treatment was performed in duplicate. On Day 3, 20 L4 larvae of each strain from each plate were transferred to fresh plates. Infection rate was assessed in the remaining Day 3 animals (F1 generation) and in all animals on Day 5 (F2 generation) by examining 50 L4 larvae or adult animals per sample using virus-specific FISH probes and protocols as described above.

Differences in infection rates across virus dilutions between each pair of wild *C. briggsae* strain and its *Cbr-gtnt-1*-edited derivative (JU1498-MCP833 and JU1564-MCP919) were evaluated for each virus and generation using the raw counts of positive and negative animals examined in FISH assays. Dilutions at which the total number of positive animals across all replicates of both strains in a pair was zero were excluded prior to analysis. For statistical methods, first, a full generalized linear model with quasi-binomial distribution, *glm(cbind(positive_n, negative_n) ∼ genotype + factor(dilution fold))*, and a reduced model excluding the “genotype” term were compared with an F-test using the R function *anova(test = “F”)*. The genotype effect was quantified as an odds ratio (OR), calculated by exponentiating the regression coefficient. This OR represents the multiplicative change in the odds of infection for one edited genotype relative to the wild-type genotype, adjusted for virus dilution. Second, for comparisons where near-complete separation was detected in the F-test (under MELV infection: JU1498-MCP833 F2, JU1564-MCP919 F1 and F2), Firth’s penalized logistic regression (via the *logistf()* function in the R package *logistif* (v1.26.1)) with binomial distribution was used instead in the two models before they were compared using the R function *anova()*. Resulting *p*-values were adjusted for multiple testing using the FDR method as above (Figure 4d, Table S4).

## Results

### Laboratory crosses using the *C. elegans* MY10 strain point to a QTL on chromosome V

The overall design of the genetic analysis of the resistance in the MY10 strain is schematized in Figure 2a. We performed a first cross between two wild *C. elegans* strains carrying the *drh-1* deletion yet differing greatly in viral load in our assay (Ashe et al. 2013): the JU1580 strain in which the Orsay virus was originally discovered (Félix et al. 2011) and the resistant MY10 strain. We crossed these two strain backgrounds and created recombinant inbred lines (see Methods) (Figure 2b). We phenotyped ORV infection in these lines using as a readout the proportion of positive animals after FISH staining of the virus. For brevity, we here sometimes refer to a low viral load or low proportion of infected individuals as viral resistance. We then sequenced resistant and sensitive pools of RILs and determined the proportion of parental alleles in each pool along the six chromosomes (Figure 2b,c). The analysis of deviations between the two pools mapped the viral resistance to a major locus on chromosome V (Figure 2c). The QTL interval started at 7.675 Mb, with the peak located left of 11 Mb and the log odds ratio decreasing greatly after 17 Mb.

To confirm the chromosome V QTL, we built chromosome substitution lines where chromosome V from one parental strain was introgressed into the background of the other parental strain and assayed viral infection using the *lys-3p::GFP* transgene as a marker (Le Pen et al. 2018). (Figure 2d). The JU4322 line carrying MY10 alleles on chromosome V in an otherwise JU1580 background was resistant to ORV infection. Conversely, the JU4323 line carrying JU1580 alleles on chromosome V in an otherwise MY10 background showed a high rate of infection, similar to the JU1580 background parent. Through screening of spontaneous recombinants, we restricted the QTL interval to 8.1-14.1 Mb (Figure 2d). We concluded that the chromosome V QTL was both necessary and sufficient to explain the resistance of MY10 when crossed into the JU1580 background.

### A non-synonymous reduction-of-function in the *gtnt-1* gene causes viral resistance of the MY10 strain

The chromosome V QTL region carries a large number of polymorphisms between JU1580 and MY10. To narrow down possible candidates, we leveraged the fact that this central region of chromosome V was recently swept in many *C. elegans* strains from across the world, including the N2 reference and the MY10 strain (Andersen et al. 2012), while the JU1580 strain (from France) instead carries a divergent chromosome V. Given the outlier resistance phenotype of MY10 in strains tested in (Ashe et al. 2013), we wondered whether the chromosome V from MY10 also caused resistance in the background of a strain with a swept chromosome V. To test this hypothesis, we chose JU1395, a wild strain closely related to the N2 reference (Andersen et al. 2012) and assayed a chromosome V introgression from MY10 into this background. Because the JU1395 genome includes an intact *drh-1* gene on chromosome IV (Ashe et al. 2013), we here assayed viral infection with the more sensitive *pals-5p::GFP* reporter, activated downstream of *drh-1* (Bakowski et al. 2014; Sowa et al. 2020). The chromosome V of MY10 introgressed in the JU1395 background did cause viral resistance (Figure 2e).

We thus narrowed down the list of candidate polymorphisms to those that, among the strains tested in (Ashe et al. 2013), were only found in the MY10 isotype at CaeNDR (Andersen et al. 2012; Cook et al. 2017) - and not in JU1395 and N2. This restricted the list to 44 polymorphisms and we did not find any large structural variant (Table S5). These 44 polymorphisms did not include frameshifts or stop codons, but included several non-synonymous alleles. After testing two candidates without success (*tofu-1* and *haao-1*), we further induced recombination for two successive rounds at 12.008 and 9.998 Mb, biasing recombination events by germline injection of the Cas9 protein and a guide RNA matching a desired location (Zdraljevic et al. 2023). Viral infection tests of the resulting recombinant lines restricted the QTL to the interval in between these two recombination breakpoints (Figure 2f).

In this 9.998-12.008 Mb interval, we tested using genome editing in the JU1580 background the non-synonymous alleles in the genes *hmp-1* (encoding an alpha-catenin gene involved in the actin cytoskeleton (Costa et al. 1998)) and *C08B6.3* (a putative glycosyltransferase; renamed *gtnt-1*, see below). We found that editing the *gtnt-1* MY10 allele (Proline to Leucine P182L substitution) resulted in viral resistance in the JU1580 background (Figure 2g) and in the reference N2 background (Figure 2h). We produced a frameshifting mutation *mf208* (which affect both predicted isoforms) in the N2 background. The resulting strain JU4465 had no overt growth or locomotion phenotype and also displayed a strong reduction in ORV infection (Figure 2i). We thus concluded that the P182L substitution in the *gtnt-1* gene caused viral resistance of the MY10 strain and is a reduction-of-function allele.

The gene encodes a putative Glycosyltransferase family 92 (GT92) protein, so we renamed it *gtnt-1* (for GlycosylTransferaseNinetyTwo92 related). Among *C. elegans* wild strains, the *gtnt-1(P182L)* allele is only present in few strains at CaeNDR (Crombie et al. 2024). However, some other wild strains carry putative loss-of-function alleles in this gene (frameshift or stop codon) (Figure 2j). We tested three such strains from distinct continents carrying three distinct mutations (Table S1) and found that they were all resistant to infection (no detectable virus by FISH; Figure 2k). EG4725 was also tested in (Ashe et al. 2013) using qRT-PCR and consistently displayed a low viral load. Thus, repeated evolution of reduction-of-function alleles occurred in *C. elegans gtnt-1*.

Expression of the *gtnt-1* gene is enriched in the intestinal cells during post-embryonic stages (Spencer et al. 2011; Cao et al. 2017; Han et al. 2017), consistent with an effect on ORV infection in these cells. *gtnt-1* expression increases in late larval stages (Wilson et al. 2023) and was not significantly altered upon ORV infection (Sarkies et al. 2013; Chen et al. 2017). We wondered whether this polymorphism affected the interaction with other intestinal pathogens. Because the transcriptional responses to viral and microsporidian infections are similar (Bakowski et al. 2014; Gonzalez and Félix 2024), we tested whether *gtnt-1* mutation altered infection by microsporidia and found that the microsporidian spore count was similar to the control (Figure 3a). Another Glycosyltransferase family 92 member, GALT-1, was previously found to be involved in resistance to the fungal lectin CGL2 (Titz et al. 2009; Butschi et al. 2010). We thus also tested whether the *gtnt-1* mutation rendered the animals resistant to nematotoxic fungal lectins and found no effect (Figure 3b). Conversely, the *galt-1* mutant displayed resistance to the CGL2 lectin (Figure 3b) but no effect on ORV infection (Figure 3c). Thus, we concluded that the pathogen resistance of the *gtnt-1* reduction-of-function allele was specific to ORV infection and did not affect microsporidian infection nor fungal lectin toxicity.

**Figure 3.**
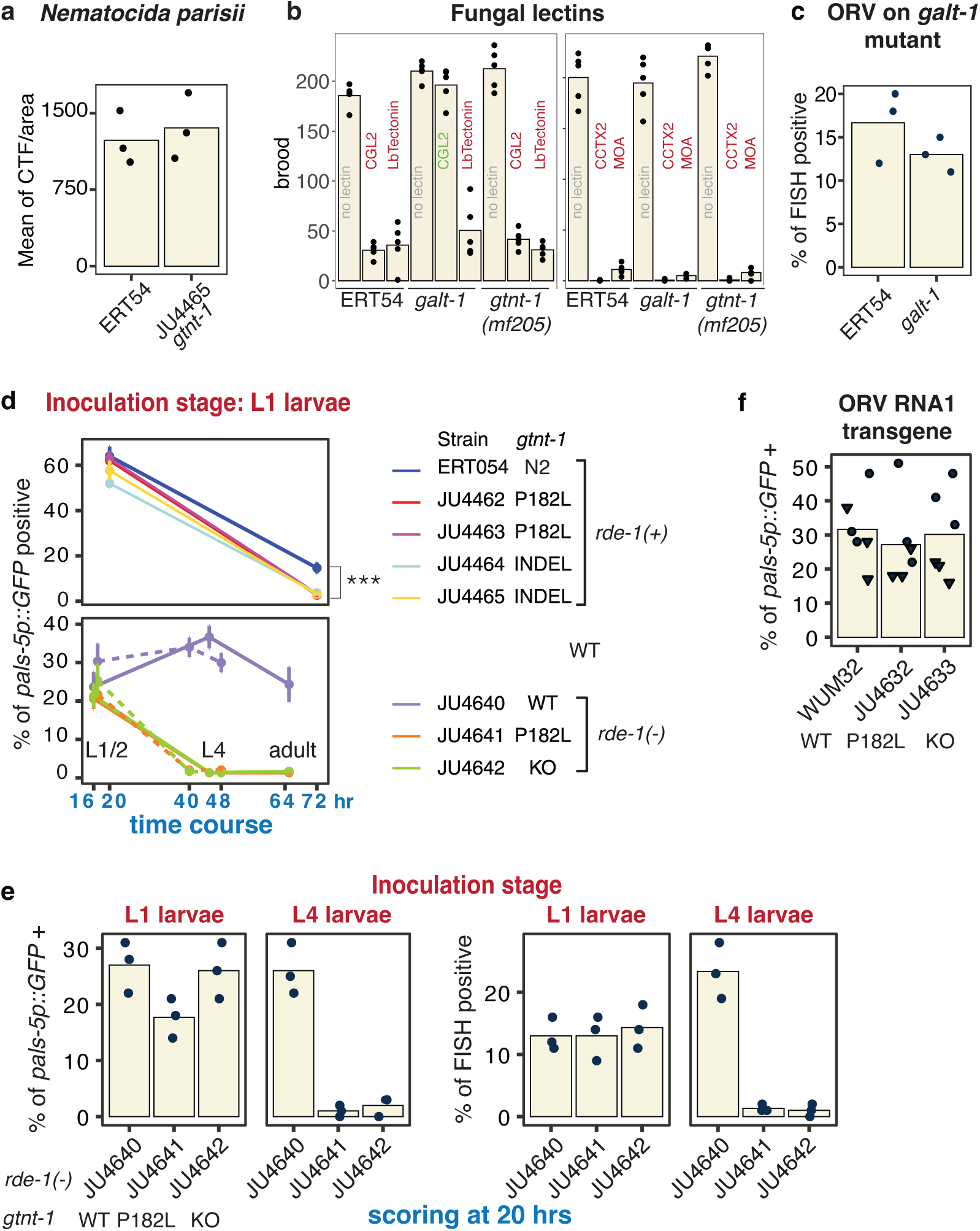
*gtnt-1* reduction-of-function mutations confer pathogen and stage-specific resistance in *C. elegans*. **a,** *gtnt-1* mutation does not confer resistance to infection by the microsporidia *Nematocida parisii* compared to wild type. Each point is a replicate showing the mean corrected total fluorescence (CTF) normalized by worm area in 60 animals. **b**, *gtnt-1* mutation does not confer resistance to five tested fungal lectins. Five fungal lectins are tested on the ERT54 control, a mutant in *galt-1*, another GT92 protein-coding gene that is resistant to the CGL2 lectin and *gtnt-1(mf205)* (strain JU4462), with five replicates per condition. **c**, The *galt-1*(*op497*) mutant is not resistant to the ORV, as assayed using FISH. **d,** Time-course measurement of infection-activated reporter (*pals-5p::GFP*) in eight *C. elegans* strains of different genetic backgrounds and *gtnt-1* genotypes (detailed on the right). Animals were infected shortly after hatching and sampled from L1/2 larval stages (20 hrs after hatching) to adults (72 hrs). Three independent assays are shown. Each dot represents the mean percentage of reporter-positive animals across three biological replicates; error bars indicate standard errors. The MY10 *gtnt-1* allele does not prevent the high activation of the reporter at early timepoints but causes a significant lower infection rate at 72 hrs compared to the reference allele (*p*<10^-11^, generalized linear model, binomial family and logit link function, comparing the *gtnt-1* N2 allele to reduction-of-function alleles). **e,** *gtnt-1* reduction-of-function mutations conferred resistance when infection occurred at the L4 stage but not at the L1 stage. Two parallel assays of viral infection are shown: infection-activated reporter (*pals-5p::GFP*) on the left and FISH-based infection quantification on the right, assayed 20 hrs post-inoculation. Each point represents one biological replicate in **d** and **e**. **f,** Heat-shock induction of a transgene expressing self-replicating ORV RNA1 carrying the viral RNA-dependent RNA polymerase, in two independent experimental blocks, indicated by dot color. A control without heat-shock shows no activation of the *pals-5p::GFP* reporter (1/100 animals scored positive in one of three replicates; Table S4).

As mentioned, our initial assays covered several days to account for the full course of viral transmission across animals. We then tested whether the resistant *gtnt-1* allele prevented entry of the virus by assaying early timepoints during infection. We inoculated the virus on plates carrying L1 stage animals and tested infection 20 and 72 hrs later. Contrary to the hypothesis of a block in viral entry, we could detect the same level of *pals-5p::GFP* activation at 20 hrs in N2 background animals carrying either *gtnt-1* allele (Figure 3d). By contrast, animals with mutations that block early steps of the viral cycle did not activate *pals-5p::GFP* reporter upon inoculation in these conditions (Jiang et al. 2017). At later timepoints (72 hrs), infection was lost in the strain carrying the *gtnt-1(P182L)*, deletion, or frameshift alleles (Figure 3d top panel). In similar time-course infection assays in a *rde-1* mutant background, wild-type *gtnt-1* also exhibited higher infection rate than those with *gtnt-1* mutations (Figure 3d bottom panel). In another experiment, we infected in parallel L1 and L4 stage animals and assayed them 20 hrs post-inoculation using *pals-5p::GFP* and ORV FISH. Again, the young larvae showed reporter activation and viral entry, while the L4 larvae with *gtnt-1* mutated alleles were resistant compared to those carrying the reference allele (Figure 3e). We concluded that *gtnt-1* loss-of-function protected the animals at late developmental stages.

We further tested whether the mutation prevented viral replication by using a strain carrying a transgene with the ORV RNA1 coding for the RNA-dependent RNA polymerase, transcribed under a heat-shock promoter in the background of the *rde-1* mutation (Jiang et al. 2017). After heat-shock, strains carrying either the N2 or the MY10 *gtnt-1* allele were able to similarly activate the *pals-5p::GF*P reporter (Figure 3f). Thus, we concluded that the *gtnt-1* mutation did not affect viral replication.

### Natural polymorphisms in *Cbr*-*gtnt-1* also confer viral resistance in *C. briggsae*

Among the 40 wild *C. briggsae* strains previously assayed for LEBV and SANTV infection (Alkan et al. 2024), ten strains (Figure 1b) showed high genetic concordance in CaeNDR (> 0.9997 of the June 2025 SNP set) and were grouped into the same isotype, NIC174 (Crombie et al. 2024). Despite their nearly identical genomes, these strains differed in infection rate upon LEBV and SANTV inoculation (Figure 1b). For instance, JU1498 and JU1264, the original strains from which LEBV and SANTV were identified, respectively (Franz et al. 2012), showed infection by both viruses (30-40% infected animals by FISH), whereas the strain JU516 appeared to displayed SANTV-specific resistance (Figure 1b) (Alkan et al. 2024).

Here, we first asked whether their different viral susceptibility led to fitness differences (Figure 4a, Table S4). When the three strains competed under the uninfected control condition, they maintained similar frequencies in the mixed population over generations. Under infection with any of the three viruses (LEBV, SANTV and MELV), the strain JU516 outcompeted the other two strains (Figure 4a), demonstrating its selective advantage, possibly resulting from resistance or tolerance to these infections.

**Figure 4.**
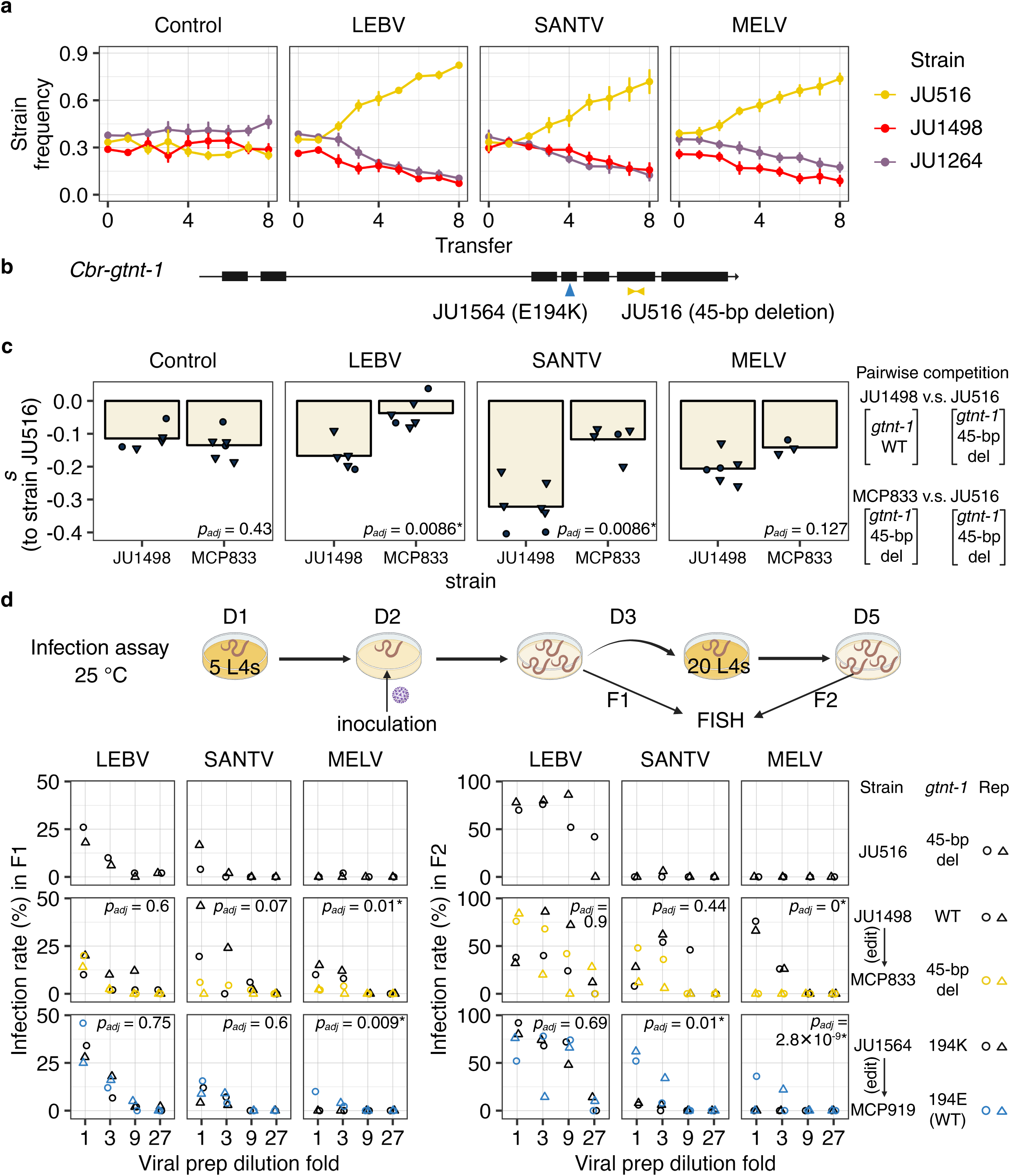
Natural variation in *Cbr*-*gtnt-1* confers virus-specific resistance in *C. briggsae*. **a,** Experimental competition among three *C. briggsae* strains of the isotype NIC174 under four conditions (uninfected control; infected with LEBV, SANTV, or MELV). Points show mean strain frequencies (*y*-axis) across 4-8 replicates (biological and technical, see Methods) at each transfer (*x*-axis); error bars indicate standard errors. Gold, red, and purple denote strains JU516, JU1498, and JU1264, respectively. **b.** Two natural variants in the gene *Cbr-gtnt-1*. **c.** Selection coefficient (y-axis) of the strains JU1498 and MCP833 relative to the strain JU516 in multigenerational pairwise competitions (approximately nine generations, see Methods). Each dot / triangle represents one biological replicate in two independent assays. Significance values (lower right in each panel) for differences in between JU1498 and MCP833 under each condition were calculated by a Wilcoxon test and were adjusted using FDR (See Methods). **d.** Infection assays of *C. briggsae* using serial dilutions of viral preparations. The scheme of the infection assay was shown above (see Methods). Three wild strains (black hollow triangles or circles) and their two *Cbr-gtnt-1* edited derivatives (colored hollow triangles or circles) were tested in parallel, each with two biological replicates. Infection rate was measured by examining ∼50 L4/adult stage animals using FISH as previously described (Frézal et al. 2019; Alkan et al. 2024). Adjusted significance values were shown upper right in each panel when applicable (see Methods). See Table S4 for raw data and statistics.

By examination of genomic variation among the *C. briggsae* JU516, JU1498, and JU1264 strains (Table S6), we found that JU516 carried a unique 45-bp deletion in the sixth exon of *CBG09519*, the *C. briggsae* ortholog of *C. elegans gtnt-1* (hereafter *Cbr-gtnt-1*) (Figure 4b, Table S6) (Sternberg et al. 2024). To test the effect of this deletion, we introduced it into *C. briggsae* JU1498 using CRISPR/Cas9 genome editing, generating the derivative MCP833. Then, we competed the two strains individually with JU516 for nine generations with or without viral infections. We compared the selection coefficient of JU1498 and MCP833 relative to JU516 between the first and the last generations (Figure 4c, Table S4). Both strains exhibited negative selection coefficients in all conditions, indicating lower fitness relative to JU516. However, the edited MCP833 showed a significantly higher selection coefficient than JU1498 under LEBV and SANTV infection (*p_adj_* = 0.0086 for both and effect size *r* = 0.83 and 0.82, respectively) and a similar trend under MELV infection (*p_adj_* = 0.13, *r* = 0.60), in contrast to their close selection coefficient under the uninfected control condition (*p_adj_* = 0.43) (Figure 4c). The JU1498 strain with the intact *Cbr-gtnt-1* was particularly affected by SANTV infection compared to control conditions. These results demonstrate that the 45-bp deletion in *Cbr-gtnt-1* enhances relative fitness of the host under viral infection.

Among the 40 strains previously assayed (Alkan et al. 2024), the *C. briggsae* strain JU1564 (isotype JU1564) was another SANTV-specific resistant strain when assayed by FISH and carries a missense variant (E194K) in *Cbr-gtnt-1* (Figure 4b) (Crombie et al. 2024). An edited derivative, MCP919, carrying the reference 194E allele, was generated in the JU1564 background to assess effects of different *Cbr-gtnt-1* natural variants on viral susceptibility of *C. briggsae*. We compared the infection rate of JU516, JU1498, and JU1564, and their two derivatives (MCP833 and MCP919) over two generations (F1 and F2) in infection assays inoculated with serially diluted viral preparations to control for differential viral preparation infectivity (Figure 4d, Table S4). Consistent with previous results (Figure 1b) (Alkan et al. 2024), JU516 and JU1564, compared with JU1498 infection at the same titrations, showed a relatively specific high infection rate with LEBV and resistance to the closely related SANTV and MELV (Figure 4d). We then compared JU1498 and JU1564 with their *Cbr-gtnt-1* edited derivatives mimicking the JU516 and JU1498 allele, respectively. For LEBV, we found no significant effect of the *Cbr-gtnt-1* genotype in either pair or generation (*p_adj_* ≥ 0.6, OR = 0.7-1.1). For SANTV, significant genotype effects were detected in JU1498-MCP833 F1 (*p* = 0.035, *p_adj_* = 0.071, OR = 0.16), and in JU1564-MCP919 F2 (*p_adj_* = 0.01, OR = 14.4), while other comparisons were not significant *(p_adj_* ≥ 0.44). For MELV, significant genotype effects were observed in JU1498-MCP833 F1 (*p_adj_* = 0.01, OR = 0.17). In other three MELV combinations, notably, infections were only detected in JU1498 and MCP919, resulting in highly significant differences *(p_adj_* ≤ 0.01). In significant cases, as expected, the allelic effects were directionally opposite: the 45-bp deletion in MCP833 decreased viral susceptibility, whereas the 194E allele in MCP919 increased susceptibility (Figure 4d).

Altogether, these results demonstrate that natural variation in *Cbr-gtnt-1* contributes to virus-specific differences in host susceptibility and fitness in *C. briggsae*.

### Repeated mutations of *gtnt-1* in two *Caenorhabditis* species

The main predicted isoform of the *gtnt-1* gene, called *gtnt-1a*, encodes a 439 amino-acid protein with a N-terminal trans-membrane (TM) domain and a Glycosyltransferase family 92 (GT92) protein domain (Figure 5a). The gene has many paralogs (Davis et al. 2022). Some GT92 members, initially found in *C. elegans*, were shown to form N-glycosylation 1,4-linkages of galactose, including in plants (Titz et al. 2009; Butschi et al. 2010; Liwanag et al. 2012; Ebert et al. 2018). GTNT-1 carries a predicted signal peptide and trans-membrane domain and a likely Golgi localization (DeepLoc2.0 (Thumuluri et al. 2022)), consistent with a possible glycosylation activity in this compartment. The polymorphism alters proline 182 (proline 3 of the short annotated GTNT-1b form) into a leucine. This amino-acid is located at the beginning of the Glycosyltransferase family 92 domain (annotated from 174) (Figure 5b). The proline is conserved in the ortholog in many *Caenorhabditis* species and thus appears to be the ancestral state. It is mutated to an asparagine in a clade formed by *C. briggsae*, *C. nigoni*, and *C. sinica* and to a threonine in *C. uteleia* (Figure 5b) (Wormbase (Davis et al. 2022)). It is also conserved in most paralogs (Davis et al. 2022).

**Figure 5.**
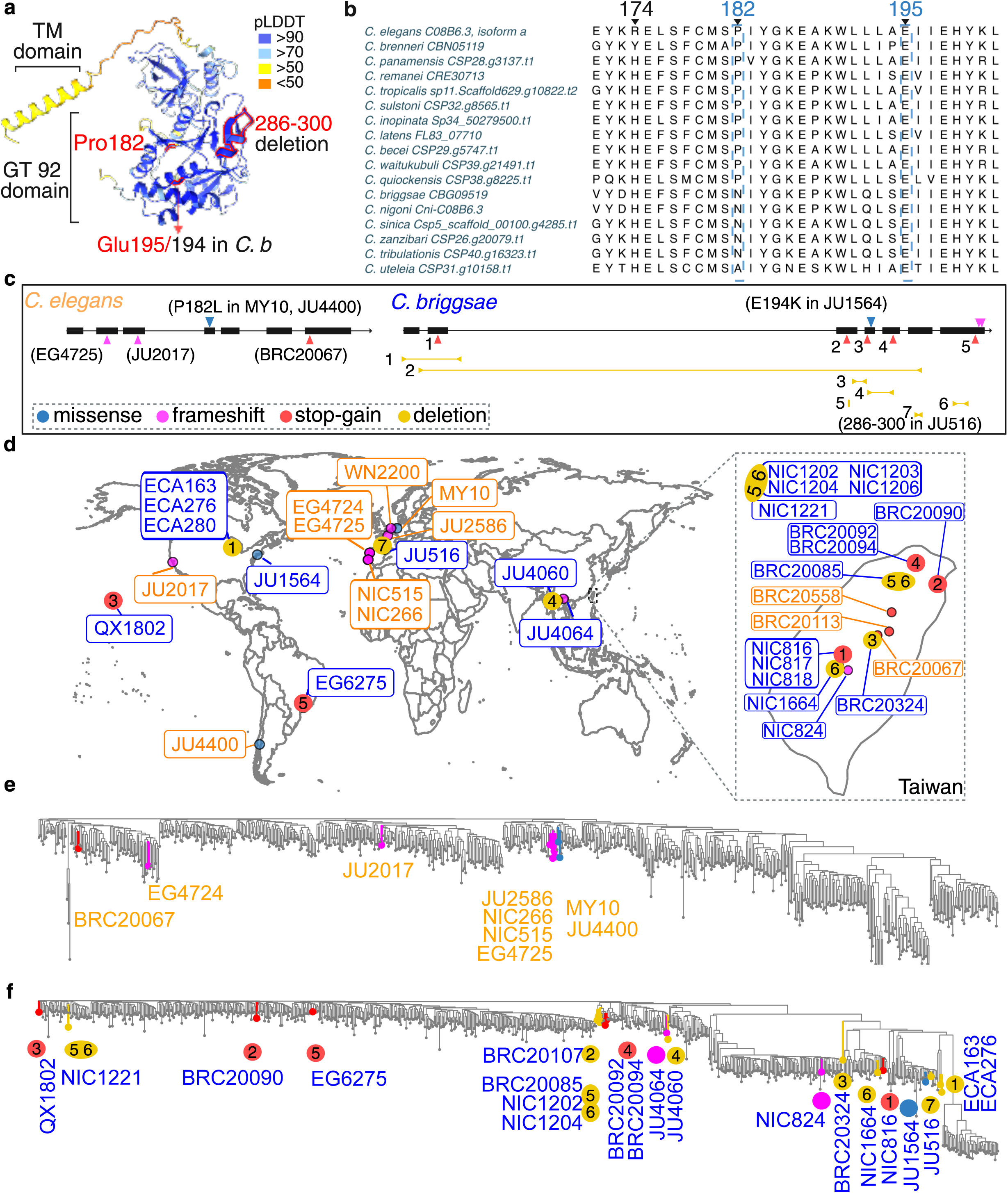
Recurrent *gtnt-1* mutations in wild *C. elegans* and C*. briggsae*. **a,** AlphaFold-predicted three-dimensional structure of GTNT-1 (Abramson et al. 2024), colored by per-residue confidence (pLDDT). The transmembrane (TM) domain and the Glycosyltransferase 92 (GT92) domain are indicated. The three experimentally validated natural mutations, P182, E195/194, and the 286-300 deletion, are outlined in red. **b,** Partial protein alignment of the *gtnt-1* orthologs in *Caenorhabditis*, using *Cel-gtnt-1* as the reference. The GT92 domain is predicted to begin at amino acid 174. **c,** Summary of variants in *gtnt-1* and its ortholog in wild *C. elegans* and *C. briggsae* strains (Crombie et al. 2024), including the two validated missense mutations (blue), frameshifts (magenta), stop-gains (red), and deletions (gold). Stop-gain and deletion variants in the *Cbr-gtnt-1* ortholog are numbered. **d,** Geographical distribution of wild *C. elegans* (orange) and *C. briggsae* (blue) strains carrying the mutations indicated in **c**. Colored points show sample locations and mutation types (numbered when applicable). **e, f,** Genetic relatedness trees of 687 wild *C. elegans* isotype strains (**e**) and 719 wild *C. briggsae* strains (**f**) from the latest CaeNDR releases, modified to highlight strains carrying the mutations indicated in **c** and mutation types.

In addition to the strain MY10 (isotype MY10), another strain from Chile, JU4400 (isotype JU4400), carries the 182L allele (Figure 5c,d) in a similar genomic background (Figure 5e), pointing to a single event of mutation to 182L. As tested in Figure 2k, several other *C. elegans* wild strains carry premature stop codons or indels and were also found to be resistant to ORV. These various alleles correspond to recurrent events of *gtnt-1* mutation forming multiple rare alleles in the species (Figure 5e).

The *C. briggsae* strain JU1564 carries a lysine at the position 194 (195 in *C. elegans*), where other *Caenorhabditis* orthologs have a conserved glutamic acid (Figure 5b). The 45-bp deletion in the *C. briggsae* strain JU516 is predicted to remove amino acids 286-300 (Figures 4b, 5a). Additionally, more natural variants in *Cbr-gtnt-1* were identified in genomes of *C. briggsae* wild strains, including multiple frameshift mutations, premature stop codons, and large deletions (Figure 5c). These mutations likely lead to reduction-of-function effects.

Strains with alternative *Cel-gtnt-1* and *Cbr-gtnt-1* alleles were isolated from across the world, with a hotspot in the main island of Taiwan (Figure 5d). They are distributed across their respective whole-genome SNV-based relatedness tree (Figure 5e, f), indicating independent origins and repeated mutations at the *gtnt-1* orthologs in both species.

## Discussion

### Natural variation reveals a host-virus interaction factor

Prior laboratory studies identified many factors involved in the viral cycle and in host immune defense using the *C. elegans*-ORV system and forward genetic screens (Jiang et al. 2017; Tanguy et al. 2017; Le Pen et al. 2018; Jiang et al. 2020; Casorla-Perez et al. 2022; Cubillas et al. 2023). The *gtnt-1* resistance alleles appear as reduction-of-function and putative null alleles without a strong lethality. In principle, their isolation would not have required particular precautions in genetic screens. Why, then, was the *gtnt-1* gene not identified? The explanation likely lies in the design of infection assays: most previous laboratory screens relied on within-generation activation of *pals-5p::GFP* transgene in the *rde-1* sensitized N2 reference background (Jiang et al. 2017; Jiang et al. 2020; Cubillas et al. 2023). Here we showed that the MY10 *gtnt-1* allele did not initially prevent *pals-5p::GFP* transgene activation in similar conditions, but prevented further spread in the population (Figure 3), perhaps corresponding to the waves of infection in (Castiglioni et al. 2024; Castiglioni et al. 2025). Therefore, our multi-generational infection assay seems to have been key in detecting the resistance of the MY10 strain population and the corresponding *gtnt-1* polymorphism. More broadly, our study also illustrates a general workflow for dissecting a QTL down to its causal variant (Figure 2a), combining RILs and pool-seq to define a genomic interval, followed by fine-mapping through spontaneous and CRISPR-Cas9-mediated recombination, and ultimately validated causality by precise genome editing at single-nucleotide resolution.

### Glycosylation in the viral lifecycle

The *gtnt-1* gene likely encodes an enzyme predicted to be involved in glycosylation. Other GT92 proteins have been biochemically demonstrated to transfer galactose in protein N-linked glycosylation (Ebert et al. 2018). Complex surface glycans constitute, in non-vertebrates but not only, a system of recognition of self and non-self, which can be perverted by pathogens (Alves et al. 2022). Lectins are proteins that recognize such specific glycosylation moieties. The glycocalyx can also simply form a physical barrier to infections (Stonebraker et al. 2004).

In *C. elegans*, genes coding for putative GT92 glycosyltransferases have been involved in resistance to various toxins and pathogens. The *Coprinopsis cinerea* fungus emits the nematotoxic lectin CGL2, and mutation in *C. elegans galt-1* renders the nematode resistant to this fungal toxin (Titz et al. 2009; Butschi et al. 2010). *Yersinia* bacteria form biofilms on *C. elegans* cuticle and mutants in the GT92 family member BAH-1 are resistant (Drace et al. 2009). Attachment of other pathogenic bacterial species (*Microbacterium nematophilum* and a *Leucobacter* species) to the *C. elegan*s cuticle is also sensitive to mutation in GT92 gene family members, such as SRF-2 and BAH-1 (O’Rourke et al. 2023). In the case of SRF-2, the lack of glycosylation presumably prevents binding of the pathogen to the cuticle.

Here we implicate glycosylation in *Caenorhabditis*-virus interactions, a novel but not surprising finding as glycosylation may affect many steps of viral cycles (Dugan et al. 2022), including for example both glycosylation of SARS-CoV-2 and its receptor ACE2 (Isobe et al. 2022). Glycosylation of viral proteins may affect their interaction with the host immune system, their folding and consequently the virus life cycle and pathogenesis (Feng et al. 2022). We note that the viral RNAs appear to be associated with endoplasmic reticulum membranes (Efstathiou et al. 2022), hence some viral proteins may be associated with intracellular membrane-bound compartments. The RdRp encoded by ORV RNA1 includes a putative signal peptide and possible Golgi-to-ER retrieval (as predicted using DeepLoc2.0 (Thumuluri et al. 2022)).

In the case of the Orsay virus, a puzzle is that the GTNT-1-dependent process does not hamper entry of the virus in early larval stages but appears to prevent entry at later stages. A possibility is that the intestinal surface matures along larval stages and that young gut cells of *gtnt-1* mutants allow for viral entry, while entry is blocked at later stages during establishment of abnormal glycosylation moieties in more mature animals. An alternative scenario is that host GTNT-1 is required at a late stage of the viral lifecycle, for example for glycosylation of a viral protein. Our experiments showing a lack of infection in short timepoints after inoculation in the L4 stage are less consistent with this second scenario. Altogether, these results suggest that the mutation either prevents viral entry in late developmental stages or act in viral maturation. Future work will address the enzymatic activity of GTNT-1 and its involvement part in the viral lifecycle.

### Recurrent mutation of *gtnt-1* leads to viral resistance in two species

Repeated loss-of-function mutations at the same gene have been proposed to provide a hotspot locus for rapid adaptation (Martin and Orgogozo 2013), which has been found in cases of resistance to toxic compounds in a diversity of animals and plants (e.g. (Karasov et al. 2010; Poormohammad Kiani et al. 2012; Hahnel et al. 2018)). In the present case, diverse *gtnt-1* reduction-of-function mutations are even found across two species, *C. elegans* and *C. briggsae.* These two species are not sister species (Kiontke et al. 2011; Stevens et al. 2019) and their reduction-of-function alleles are not shared between the two species, which appears to rule out a common origin of alleles. We thus discovered within *Caenorhabditis* a clear case of repeated evolution (Martin and Orgogozo 2013; Cerca 2023), where the *gtnt-1* gene is a hotspot of derived mutations for viral resistance in two species that are infected by distinct viruses of the same family. An ancestral virus might co-opt *gtnt-1*-dependent host functions, and this dependency was retained as host and virus lineages diverged, making it a recurrent target of selection. The resistant *gtnt-1* alleles are found in host populations across the world. The geographic distribution of the viruses is not well known as sampling is heavily biased by the location of our laboratory in Europe (Frézal et al. 2019). The *gtnt-1* locus is not the only source of variation in host-virus interaction in either species (Ashe et al. 2013; Alkan et al. 2024), however it is the only one known so far to have mutated towards resistance - in contrast to the derived intermediate-frequency *C. elegans drh-1* deletion which promotes sensitivity (Ashe et al. 2013).

The variable environment conferred by pathogens is thought to favor the maintenance of host immune system polymorphisms in natural populations. The genomes of both *C. elegans* and *C. briggsae* contain hyperdivergent regions, which are enriched with immune response genes and have been hypothesized to be maintained at intermediate frequency by balancing selection such as that operated by pathogens (Lee et al. 2021). In striking contrast, the viral resistant alleles of *gtnt-1* are recent mutations, each occurring at low frequency (<1% of sequenced strains and isotypes). Orthologs of *gtnt-1* are found throughout *Caenorhabditis* (Figure 5), thus, despite recurrent mutation to putative null alleles (nonsense, indels), the gene does not degenerate in the longer evolutionary term. That these loss-of-function alleles are neutral in natural populations remains a possibility, however they cause increase fitness in laboratory conditions in the presence of the viruses (Ashe et al. 2013; Alkan et al. 2024) (the present work). Possible explanations for their low frequency are the following: first, *gtnt-1* may be pleiotropic and required for fitness-related phenotypes in other environments, in which case, its degeneration may be transiently advantageous in the face of viral infections but deleterious in the long term. Second, a related explanation could be that the viral sensitivity itself may be advantageous in some conditions, such as enabling activation of host responses that protect against stronger pathogens, for example the intracellular pathogen response protecting against microsporidia or heat shock (Reddy et al. 2017; Reddy et al. 2019; Castiglioni and Elena 2024). In this case, the ancestral allele may never be lost despite viral sensitivity and loss of fitness in some conditions. A similar case is well known in the regulation of flowering in *Arabidopsis*, with repeated loss-of-function alleles in the *FRIGIDA* gene regulating flowering in relation with winter temperature (Johanson et al. 2000; Le Corre et al. 2002; Zhang and Jimenez-Gomez 2020). Mutation at the locus does not seem limiting the evolution of resistance (or flowering in warm conditions), resulting in high allelic heterogeneity.

The effects of loss-of-function *gtnt-1* alleles varied across specific host-virus interactions. Similar to the responses of *C. elegans* to ORV, *Cbr-gtnt-1* polymorphisms significantly reduced the proportion of infected *C. briggsae* upon SANTV and MELV but not LEBV (Figure 4d). This exception of LEBV likely reflects the phylogenetic relationships among the four *Caenorhabditis* viruses: LEBV is the most divergent in their RdRp phylogeny, forming the outgroup to a clade comprising ORV, SANTV, and MELV, within which the latter two clustered together (Frézal et al. 2019). However, the *Cbr-gtnt-1* reduction-, or loss-of-function allele appeared to increase host relative fitness in the presence of all three viruses (Figure 4c), suggesting that it also affected infection dynamics and/or tolerance to LEBV. LEBV may have evolved to only partially depend on *Cbr-gtnt-1*, allowing it to maintain infection rates under reduction-of-function *Cbr-gtnt-1*. The virulence of LEBV may ease without *Cbr-gtnt-1*, such as reduced viral loads per host cell, which could improve host relative fitness. These differences among the host-virus pairs suggest that closely related viruses can evolve distinct dependencies on the same host factor, and dissecting the molecular basis of this variation will help clarify which steps of the viral lifecycle are affected and how these interactions have diverged in host-virus co-evolution.

## Conclusion

We found repeated evolution of reduction-of-function alleles of *gtnt-1* orthologs in two nematode species that confer viral resistance but remain at low frequency. These results highlight how mutations in a single gene can provide repeatable solutions to adaptation across species and populations.

## Acknowledgements

We thank Lise Frézal for help with the initial genomic analyses, Jonathan Hodgkin, Iain Wilson and Markus Kuenzler for discussions and sending the fungal lectin plasmids. We thank Deborah Bourchis’ laboratory (Curie Institute) for access to their pyrosequencer. We thank Katie Pelletier for comments on the manuscript. This work was supported by grants from the Agence Nationale pour le Recherche ANR-11-BSV3-013 and from the Fondation pour la Recherche Médicale DEQ20150331704 to MAF and ARF202209015859 to GZ. We thank Wormbase and CaeNDR. Some strains were provided by the CGC, which is funded by NIH Office of Research Infrastructure Programs (P40 OD010440). Some strains were generated by SEGiCel (SFR Santé Lyon Est CNRS UAR 3453, Lyon, France) with the support of CNRS and IBiSA - we thank especially Arnaud Hochard and Margaux Gibert.

## Supplementary Material

**Table S1. Strain list.** This table lists in columns: strain, genotype, strain origin (reference), comments, sequence of the region of CRISPR edits in nucleotides and predicted resulting amino-acids. In the latter two columns, mutations are indicated in red font.

**Table S2. Oligonucleotides.** PCR primers, pyrosequencing primers and CRISPR oligonucleotides are indicated in successive sheets.

**Table S3. Frequency of the MY10 allele at the 46,833 single-nucleotide polymorphism markers, in the Orsay-virus resistant and sensitive *C. elegans* pools.** The first and second columns indicate the chromosome and nucleotide position, the third and fourth the proportion of reads in the resistant and sensitive pools, respectively. These data are binned and plotted in Figure 2b.

**Table S4 Infection data**. Raw data corresponding to the infection experiments in Figures 2, 3, and 4.

**Table S5. Candidate polymorphisms on chromosome V.** This table indicates MY10-specific polymorphisms in the region defined in Figure 2d. See Methods for filters. The MY10 strain carries a center of chromosome V that is very similar to that of the reference strain N2, defined as the swept haplotype in (Andersen et al. 2012). Therefore, the number of MY10-specific polymorphisms is low and they represent recent mutations. We did not find any structural variant in the region beyond the small indels of this table. Tested candidates are highlighted in bold. The final QTL region defined by recombinant mapping in Figure 2f is highlighted in light blue. The causal variant is highlighted in yellow.

**Table S6. Variants between *C. briggsae* JU516, JU1498 and JU1264, filtering for nonsynonymous, nonsense, and indel variants.**

## Notes

### Competing Interest Statement

The authors have declared no competing interest.

## References

Abramson, J., Adler, J., Dunger, J., Evans, R., et al. (2024). Accurate structure prediction of biomolecular interactions with AlphaFold 3. Nature 630: 493–500. https://www.ncbi.nlm.nih.gov/pubmed/38718835

Alkan, C., Brésard, G., Frézal, L., Richaud, A., Ruaud, A., Zhang, G., Félix, M.-A. (2024). Natural variation in infection specificity of *Caenorhabditis briggsae* isolates by two RNA viruses. PLoS Pathog 20: e1012259. https://www.ncbi.nlm.nih.gov/pubmed/38861582

Alves, I., Fernandes, A., Santos-Pereira, B., Azevedo, C. M., Pinho, S. S. (2022). Glycans as a key factor in self and nonself discrimination: impact on the breach of immune tolerance. FEBS Lett 596: 1485–1502. https://www.ncbi.nlm.nih.gov/pubmed/35383918

Andersen, E., Gerke, J., Shapiro, J., Crissman, J., Ghosh, R., Bloom, J., Félix, M.-A., Kruglyak, L. (2012). Chromosome-scale selective sweeps shape *Caenorhabditis elegans* genomic diversity. Nature Genetics 45: 285–290. http://www.ncbi.nlm.nih.gov/pubmed/22286215

Andersen, E. C., Rockman, M. V. (2022). Natural genetic variation as a tool for discovery in *Caenorhabditis* nematodes. Genetics 220: iyab156. https://www.ncbi.nlm.nih.gov/pubmed/35134197

Arribere, J. A., Bell, R. T., Fu, B. X., Artiles, K. L., Hartman, P. S., Fire, A. Z. (2014). Efficient marker-free recovery of custom genetic modifications with CRISPR/Cas9 in *Caenorhabditis elegans*. Genetics 198: 837–46. http://www.ncbi.nlm.nih.gov/pubmed/25161212

Ashe, A., Bélicard, T., Le Pen, J., Sarkies, P., Frézal, L., Lehrbach, N. J., Félix, M. A., Miska, E. A. (2013). A deletion polymorphism in the *Caenorhabditis elegans* RIG-I homolog disables viral RNA dicing and antiviral immunity. Elife 2: e00994. http://ncbi.nlm.nih.gov/pubmed/24137537

Bakowski, M. A., Desjardins, C. A., Smelkinson, M. G., Dunbar, T. A., Lopez-Moyado, I. F., Rifkin, S. A., Cuomo, C. A., Troemel, E. R. (2014). Ubiquitin-mediated response to microsporidia and virus infection in *C. elegans*. PLoS Pathog 10: e1004200. http://www.ncbi.nlm.nih.gov/pubmed/24945527

Balla, K. M., Andersen, E. C., Kruglyak, L., Troemel, E. R. (2015). A wild *C. elegans* strain has enhanced epithelial immunity to a natural microsporidian parasite. PLoS Pathog 11: e1004583. http://www.ncbi.nlm.nih.gov/pubmed/25680197

Butschi, A., Titz, A., Walti, M. A., Olieric, V., Paschinger, K., Nobauer, K., Guo, X., Seeberger, P. H., Wilson, I. B., Aebi, M., Hengartner, M. O., Kunzler, M. (2010). *Caenorhabditis elegans* N-glycan core beta-galactoside confers sensitivity towards nematotoxic fungal galectin CGL2. PLoS Pathog 6: e1000717. https://www.ncbi.nlm.nih.gov/pubmed/20062796

Cao, J., Packer, J. S., Ramani, V., Cusanovich, D. A., Huynh, C., Daza, R., Qiu, X., Lee, C., Furlan, S. N., Steemers, F. J., Adey, A., Waterston, R. H., Trapnell, C., Shendure, J. (2017). Comprehensive single-cell transcriptional profiling of a multicellular organism. Science 357: 661–667. https://www.ncbi.nlm.nih.gov/pubmed/28818938

Casorla-Perez, L. A., Guennoun, R., Cubillas, C., Peng, B., Kornfeld, K., Wang, D. (2022). Orsay Virus infection of *Caenorhabditis elegans* is modulated by zinc and dependent on lipids. J Virol 96: e0121122. https://www.ncbi.nlm.nih.gov/pubmed/36342299

Castiglioni, V. G., Elena, S. F. (2024). Orsay virus infection increases *Caenorhabditis elegans* resistance to heat-shock. Biol Lett 20: 20240278. https://www.ncbi.nlm.nih.gov/pubmed/39137892

Castiglioni, V. G., Olmo-Uceda, M. J., Villena-Gimenez, A., Munoz-Sanchez, J. C., Legarda, E. G., Elena, S. F. (2024). Story of an infection: Viral dynamics and host responses in the *Caenorhabditis elegans*-Orsay virus pathosystem. Sci Adv 10: eadn5945. https://www.ncbi.nlm.nih.gov/pubmed/39331715

Castiglioni, V. G., Villena-Gimenez, A., Herek, D., Gonzalez-Sanchez, A., Toft, C., Gomez, G. G., Elena, S. F. (2025). Latent infection of *Caenorhabditis elegans* by Orsay virus induces age-dependent immunity and cross-protection. Nat Commun 16: 7123. https://www.ncbi.nlm.nih.gov/pubmed/40753171

Cerca, J. (2023). Understanding natural selection and similarity: Convergent, parallel and repeated evolution. Mol Ecol 32: 5451–5462. https://www.ncbi.nlm.nih.gov/pubmed/37724599

Chen, K., Franz, C. J., Jiang, H., Jiang, Y., Wang, D. (2017). An evolutionarily conserved transcriptional response to viral infection in *Caenorhabditis* nematodes. BMC Genomics 18: 303. https://www.ncbi.nlm.nih.gov/pubmed/28415971

Chen, S., Zhou, Y., Chen, Y., Gu, J. (2018). fastp: an ultra-fast all-in-one FASTQ preprocessor. Bioinformatics 34: i884–i890. https://www.ncbi.nlm.nih.gov/pubmed/30423086

Cook, D. E., Zdraljevic, S., Roberts, J. P., Andersen, E. C. (2017). CeNDR, the *Caenorhabditis elegans* natural diversity resource. Nucleic Acids Res 45: D650–D657. https://www.ncbi.nlm.nih.gov/pubmed/27701074

Cordara, G., Egge-Jacobsen, W., Johansen, H. T., Winter, H. C., Goldstein, I. J., Sandvig, K., Krengel, U. (2011). *Marasmius oreades* agglutinin (MOA) is a chimerolectin with proteolytic activity. Biochem Biophys Res Commun 408: 405–10. https://www.ncbi.nlm.nih.gov/pubmed/21513701

Costa, M., Raich, W., Agbunag, C., Leung, B., Hardin, J., Priess, J. R. (1998). A putative catenin-cadherin system mediates morphogenesis of the *Caenorhabditis elegans* embryo. J. Cell Biol. 141: 297–308.

Crombie, T. A., McKeown, R., Moya, N. D., Evans, K. S., et al. (2024). CaeNDR, the *Caenorhabditis* Natural Diversity Resource. Nucleic Acids Res 52: D850–D858. https://www.ncbi.nlm.nih.gov/pubmed/37855690

Cubillas, C., Sandoval Del Prado, L. E., Goldacker, S., Fujii, C., Pinski, A. N., Zielke, J., Wang, D. (2023). The *alg-1* gene Is necessary for Orsay Virus replication in *Caenorhabditis elegans*. J Virol **97**: e0006523. https://www.ncbi.nlm.nih.gov/pubmed/37017532

Danecek, P., Bonfield, J. K., Liddle, J., Marshall, J., Ohan, V., Pollard, M. O., Whitwham, A., Keane, T., McCarthy, S. A., Davies, R. M., Li, H. (2021). Twelve years of SAMtools and BCFtools. Gigascience 10: giab008. https://www.ncbi.nlm.nih.gov/pubmed/33590861

Davis, P., Zarowiecki, M., Arnaboldi, V., Becerra, A., et al. (2022). WormBase in 2022-data, processes, and tools for analyzing *Caenorhabditis elegans*. Genetics 220: iyac003. https://www.ncbi.nlm.nih.gov/pubmed/35134929

Drace, K., McLaughlin, S., Darby, C. (2009). *Caenorhabditis elegans* BAH-1 is a DUF23 protein expressed in seam cells and required for microbial biofilm binding to the cuticle. PLoS One 4: e6741. https://www.ncbi.nlm.nih.gov/pubmed/19707590

Dubois, C., Felix, M. A. (2023). A QTL on chromosome IV explains a natural variation of QR.pap final position in Caenorhabditis elegans. MicroPubl Biol 2023. https://www.ncbi.nlm.nih.gov/pubmed/37273577

Dugan, A. E., Peiffer, A. L., Kiessling, L. L. (2022). Advances in glycoscience to understand viral infection and colonization. Nat Methods 19: 384–387. https://www.ncbi.nlm.nih.gov/pubmed/35396476

Duveau, F., Félix, M.-A. (2012). Role of pleiotropy in the evolution of a cryptic developmental variation in *C. elegans*. PLoS Biol 10: e1001230. http://ncbi.nlm.nih.gov/pubmed/22235190

Ebert, B., Birdseye, D., Liwanag, A. J. M., Laursen, T., et al. (2018). The three Members of the Arabidopsis glycosyltransferase family 92 are functional beta-1,4-galactan dynthases. Plant Cell Physiol 59: 2624–2636. https://www.ncbi.nlm.nih.gov/pubmed/30184190

Efstathiou, S., Ottens, F., Schutter, L. S., Ravanelli, S., Charmpilas, N., Gutschmidt, A., Le Pen, J., Gehring, N. H., Miska, E. A., Boucas, J., Hoppe, T. (2022). ER-associated RNA silencing promotes ER quality control. Nat Cell Biol 24: 1714–1725. https://www.ncbi.nlm.nih.gov/pubmed/36471127

Félix, M.-A., Ashe, A., Piffaretti, J., Wu, G., Nuez, I., Bélicard, T., Jiang, Y., Zhao, G., Franz, C. J., Goldstein, L. D., Sanroman, M., Miska, E. A., Wang, D. (2011). Natural and experimental infection of *Caenorhabditis* nematodes by novel viruses related to nodaviruses. PLoS Biol. 9: e1000586. https://journals.plos.org/plosbiology/article?id=10.1371/journal.pbio.1000586

Félix, M. A., Wang, D. (2019). Natural viruses of *Caenorhabditis* nematodes. Annu Rev Genet 53: 313–326. https://www.ncbi.nlm.nih.gov/pubmed/31424970

Feng, T., Zhang, J., Chen, Z., Pan, W., Chen, Z., Yan, Y., Dai, J. (2022). Glycosylation of viral proteins: Implication in virus-host interaction and virulence. Virulence 13: 670–683. https://www.ncbi.nlm.nih.gov/pubmed/35436420

Franz, C. J., Renshaw, H., Frezal, L., Jiang, Y., Félix, M. A., Wang, D. (2014). Orsay, Santeuil and Le Blanc viruses primarily infect intestinal cells in *Caenorhabditis* nematodes. Virology 448: 255–64. http://www.ncbi.nlm.nih.gov/pubmed/24314656

Franz, C. J., Zhao, G., Félix, M.-A., Wang, D. (2012). Complete genome sequence of Le Blanc virus, a third *Caenorhabditis* nematode-infecting virus. J. Virol. 86: 11940.

Frézal, L., Demoinet, E., Braendle, C., Miska, E. A., Félix, M.-A. (2018). Natural genetic variation in a multigenerational phenotype in *C. elegans*. Curr Biol 28: 2588–2596. https://pubmed.ncbi.nlm.nih.gov/30078564/

Frézal, L., Jung, H., Tahan, S., Wang, D., Félix, M.-A. (2019). Noda-like RNA viruses infecting *Caenorhabditis* nematodes: sympatry, diversity and reassortment. J. Virol. 93: e01170–19. https://journals.asm.org/doi/10.1128/jvi.01170-19

Gonzalez, R., Félix, M.-A. (2024). *Caenorhabditis elegans* immune responses to microsporidia and viruses. Dev Comp Immunol 154: 105148. https://www.ncbi.nlm.nih.gov/pubmed/38325500

Guo, X., Lu, R. (2013). Characterization of virus-encoded RNA interference suppressors in *Caenorhabditis elegans*. J Virol 87: 5414–23. https://www.ncbi.nlm.nih.gov/pubmed/23468484

Hahnel, S. R., Zdraljevic, S., Rodriguez, B. C., Zhao, Y., McGrath, P. T., Andersen, E. C. (2018). Extreme allelic heterogeneity at a *Caenorhabditis elegans* beta-tubulin locus explains natural resistance to benzimidazoles. PLoS Pathog 14: e1007226. https://www.ncbi.nlm.nih.gov/pubmed/30372484

Han, S., Schroeder, E. A., Silva-Garcia, C. G., Hebestreit, K., Mair, W. B., Brunet, A. (2017). Mono-unsaturated fatty acids link H3K4me3 modifiers to *C. elegans* lifespan. Nature 544: 185–190. https://www.ncbi.nlm.nih.gov/pubmed/28379943

Isobe, A., Arai, Y., Kuroda, D., Okumura, N., Ono, T., Ushiba, S., Nakakita, S. I., Daidoji, T., Suzuki, Y., Nakaya, T., Matsumoto, K., Watanabe, Y. (2022). ACE2 N-glycosylation modulates interactions with SARS-CoV-2 spike protein in a site-specific manner. Commun Biol 5: 1188. https://www.ncbi.nlm.nih.gov/pubmed/36335195

Jiang, H., Chen, K., Sandoval, L. E., Leung, C., Wang, D. (2017). An evolutionarily conserved pathway essential for Orsay virus infection of *Caenorhabditis elegans*. MBio 8: e00940–17. https://www.ncbi.nlm.nih.gov/pubmed/28874467

Jiang, H., Sandoval Del Prado, L. E., Leung, C., Wang, D. (2020). Huntingtin-interacting protein family members have a conserved pro-viral function from *Caenorhabditis elegans* to humans. Proc Natl Acad Sci U S A 117: 22462–22472. https://www.ncbi.nlm.nih.gov/pubmed/32839311

Johanson, U., West, J., Lister, C., Michaels, S., Amasino, R., Dean, C. (2000). Molecular analysis of *FRIGIDA*, a major determinant of natural variation in *Arabidopsis* flowering time. Science 290: 344–7. http://www.ncbi.nlm.nih.gov/pubmed/11030654

Karasov, T., Messer, P. W., Petrov, D. A. (2010). Evidence that adaptation in *Drosophila* is not limited by mutation at single sites. PLoS Genet 6: e1000924. https://www.ncbi.nlm.nih.gov/pubmed/20585551

Kiontke, K., Félix, M.-A., Ailion, M., Rockman, M. V., Braendle, C., Pénigault, J.-B., Fitch, D. H. (2011). A phylogeny and molecular barcodes for *Caenorhabditis*, with numerous new species from rotting fruits. BMC Evol. Biol. 11: 339. https://www.ncbi.nlm.nih.gov/pubmed/22103856

Le Corre, V., Roux, F., Reboud, X. (2002). DNA polymorphism at the *FRIGIDA* gene in *Arabidopsis thaliana*: extensive nonsynonymous variation is consistent with local selection for flowering time. Mol Biol Evol 19: 1261–71. https://www.ncbi.nlm.nih.gov/pubmed/12140238

Le Pen, J., Jiang, H., Di Domenico, T., Kneuss, E., Kosalka, J., Leung, C., Morgan, M., Much, C., Rudolph, K. L. M., Enright, A. J., O’Carroll, D., Wang, D., Miska, E. A. (2018). Terminal uridylyltransferases target RNA viruses as part of the innate immune system. Nat Struct Mol Biol 25: 778–786. https://www.ncbi.nlm.nih.gov/pubmed/30104661

Lee, D., Zdraljevic, S., Stevens, L., Wang, Y., et al. (2021). Balancing selection maintains hyper-divergent haplotypes in *Caenorhabditis elegans*. Nat Ecol Evol 5: 794–807. https://www.ncbi.nlm.nih.gov/pubmed/33820969

Li, H., Durbin, R. (2009). Fast and accurate short read alignment with Burrows-Wheeler transform. Bioinformatics 25: 1754–60. https://www.ncbi.nlm.nih.gov/pubmed/19451168

Liwanag, A. J., Ebert, B., Verhertbruggen, Y., Rennie, E. A., Rautengarten, C., Oikawa, A., Andersen, M. C., Clausen, M. H., Scheller, H. V. (2012). Pectin biosynthesis: GALS1 in *Arabidopsis thaliana* is a beta-1,4-galactan beta-1,4-galactosyltransferase. Plant Cell 24: 5024–36. https://www.ncbi.nlm.nih.gov/pubmed/23243126

Martin, A., Orgogozo, V. (2013). The loci of repeated evolution: a catalog of genetic hotspots of phenotypic variation. Evolution 67: 1235–50. http://www.ncbi.nlm.nih.gov/pubmed/23617905

Milne, I., Stephen, G., Bayer, M., Cock, P. J., Pritchard, L., Cardle, L., Shaw, P. D., Marshall, D. (2013). Using Tablet for visual exploration of second-generation sequencing data. Brief Bioinform 14: 193–202. https://www.ncbi.nlm.nih.gov/pubmed/22445902

O’Rourke, D., Gravato-Nobre, M. J., Stroud, D., Pritchett, E., Barker, E., Price, R. L., Robinson, S. A., Spiro, S., Kuwabara, P., Hodgkin, J. (2023). Isolation and molecular identification of nematode surface mutants with resistance to bacterial pathogens. G3 (Bethesda) 13. https://www.ncbi.nlm.nih.gov/pubmed/36911920

Poormohammad Kiani, S., Trontin, C., Andreatta, M., Simon, M., Robert, T., Salt, D. E., Loudet, O. (2012). Allelic heterogeneity and trade-off shape natural variation for response to soil micronutrient. PLoS Genet 8: e1002814. https://www.ncbi.nlm.nih.gov/pubmed/22807689

Reddy, K. C., Dror, T., Sowa, J. N., Panek, J., Chen, K., Lim, E. S., Wang, D., Troemel, E. R. (2017). An intracellular pathogen response pathway promotes proteostasis in *C. elegans*. Curr Biol 27: 3544–3553. https://www.ncbi.nlm.nih.gov/pubmed/29103937

Reddy, K. C., Dror, T., Underwood, R. S., Osman, G. A., Elder, C. R., Desjardins, C. A., Cuomo, C. A., Barkoulas, M., Troemel, E. R. (2019). Antagonistic paralogs control a switch between growth and pathogen resistance in *C. elegans*. PLoS Pathog 15: e1007528. https://www.ncbi.nlm.nih.gov/pubmed/30640956

Sarkies, P., Ashe, A., Le Pen, J., McKie, M. A., Miska, E. A. (2013). Competition between virus-derived and endogenous small RNAs regulates gene expression in *Caenorhabditis elegans*. Genome Res 23: 1258–70. http://www.ncbi.nlm.nih.gov/pubmed/23811144

Schmieder, S. S., Cordara, G., Kersten, F., Steiner, K., Samin, C. H., Plaza, D. F., Ahmad, A. A., Boeggild, A., Karlsen, J. L., Sokolowska, B. O., Boesen, T., Krengerl, U., Künzler, M. (2025). Structure and function of a fungal AB toxin-like chimerolectin involved in anti-nematode defense. bioRxiv, doi:10.1101/2025.08.11.669643 DOI: 10.1101/2025.08.11.669643.

Sowa, J. N., Jiang, H., Somasundaram, L., Tecle, E., Xu, G., Wang, D., Troemel, E. R. (2020). The *Caenorhabditis elegans* RIG-I homolog DRH-1 mediates the intracellular pathogen response upon viral infection. J Virol 94: e01173–19. https://www.ncbi.nlm.nih.gov/pubmed/31619561

Spencer, W. C., Zeller, G., Watson, J. D., Henz, S. R., et al. (2011). A spatial and temporal map of *C. elegans* gene expression. Genome Res 21: 325–41. https://www.ncbi.nlm.nih.gov/pubmed/21177967

Sternberg, P. W., Van Auken, K., Wang, Q., Wright, A., et al. (2024). WormBase 2024: status and transitioning to Alliance infrastructure. Genetics 227: iyae050. https://www.ncbi.nlm.nih.gov/pubmed/38573366

Stevens, L., Félix, M.-A., Beltran, T., Braendle, C., et al. (2019). Comparative genomics of ten new *Caenorhabditis* species. Evolution Letters 3: 217–236. https://onlinelibrary.wiley.com/doi/full/10.1002/evl3.110

Stiernagle, T. (2006). “Maintenance of *C. elegans*.” Wormbook, from https://www.ncbi.nlm.nih.gov/pubmed/18050451. DOI: doi/10.1895/wormbook.1.101.1.

Stonebraker, J. R., Wagner, D., Lefensty, R. W., Burns, K., Gendler, S. J., Bergelson, J. M., Boucher, R. C., O’Neal, W. K., Pickles, R. J. (2004). Glycocalyx restricts adenoviral vector access to apical receptors expressed on respiratory epithelium in vitro and in vivo: role for tethered mucins as barriers to lumenal infection. J Virol 78: 13755–68. https://www.ncbi.nlm.nih.gov/pubmed/15564484

Tanguy, M., Veron, L., Stempor, P., Ahringer, J., Sarkies, P., Miska, E. A. (2017). An alternative STAT signaling pathway acts in viral immunity in *Caenorhabditis elegans*. MBio 8: e00924–17. https://www.ncbi.nlm.nih.gov/pubmed/28874466

Thumuluri, V., Almagro Armenteros, J. J., Johansen, A. R., Nielsen, H., Winther, O. (2022). DeepLoc 2.0: multi-label subcellular localization prediction using protein language models. Nucleic Acids Res 50: W228–W234. https://www.ncbi.nlm.nih.gov/pubmed/35489069

Titz, A., Butschi, A., Henrissat, B., Fan, Y. Y., Hennet, T., Razzazi-Fazeli, E., Hengartner, M. O., Wilson, I. B. H., Kunzler, M., Aebi, M. (2009). Molecular basis for galactosylation of core fucose residues in invertebrates: identification of *Caenorhabditis elegans* N-glycan core alpha1,6-fucoside beta1,4-galactosyltransferase GALT-1 as a member of a novel glycosyltransferase family. J Biol Chem 284: 36223–36233. https://www.ncbi.nlm.nih.gov/pubmed/19858195

Wernet, N. D., Tecle, E., Sarmiento, M. B., Kuo, C. J., Chhan, C. B., Baick, I., Batachari, L. E., Franklin, L., Herneisen, A., Bhabha, G., Ekiert, D. C., Hanna-Rose, W., Troemel, E. R. (2025). Adenosine deaminase and deoxyadenosine regulate intracellular immune response in *C. elegans*. iScience 28: 111950. https://www.ncbi.nlm.nih.gov/pubmed/40034845

Wilson, I. B. H., Yan, S., Jin, C., Dutkiewicz, Z., Rendic, D., Palmberger, D., Schnabel, R., Paschinger, K. (2023). Increasing complexity of the N-glycome during *Caenorhabditis* development. Mol Cell Proteomics 22: 100505. https://www.ncbi.nlm.nih.gov/pubmed/36717059

Wohlschlager, T., Butschi, A., Grassi, P., Sutov, G., Gauss, R., Hauck, D., Schmieder, S. S., Knobel, M., Titz, A., Dell, A., Haslam, S. M., Hengartner, M. O., Aebi, M., Kunzler, M. (2014). Methylated glycans as conserved targets of animal and fungal innate defense. Proc Natl Acad Sci U S A 111: E2787–96. https://www.ncbi.nlm.nih.gov/pubmed/24879441

Wohlschlager, T., Butschi, A., Zurfluh, K., Vonesch, S. C., Auf dem Keller, U., Gehrig, P., Bleuler-Martinez, S., Hengartner, M. O., Aebi, M., Kunzler, M. (2011). Nematotoxicity of *Marasmius oreades* agglutinin (MOA) depends on glycolipid binding and cysteine protease activity. J Biol Chem 286: 30337–30343. https://www.ncbi.nlm.nih.gov/pubmed/21757752

Ye, K., Schulz, M. H., Long, Q., Apweiler, R., Ning, Z. (2009). Pindel: a pattern growth approach to detect break points of large deletions and medium sized insertions from paired-end short reads. Bioinformatics 25: 2865–2871.

Zdraljevic, S., Walter-McNeill, L., Marquez, H., Kruglyak, L. (2023). Heritable Cas9-induced nonhomologous recombination in *C. elegans*. MicroPubl Biol 2023 DOI: 10.17912/micropub.biology.000775.

Zhang, L., Jimenez-Gomez, J. M. (2020). Functional analysis of *FRIGIDA* using naturally occurring variation in *Arabidopsis thaliana*. Plant J 103: 154–165. https://www.ncbi.nlm.nih.gov/pubmed/32022960

